# LST-1 acts in *trans* with a conserved RNA-binding protein to maintain stem cells

**DOI:** 10.1101/657312

**Authors:** Kimberly A. Haupt, Amy L. Enright, Ahlan S. Ferdous, Aaron M. Kershner, Heaji Shin, Marvin Wickens, Judith Kimble

**Affiliations:** Department of Biochemistry, University of Wisconsin-Madison, Madison, Wisconsin, 53706, USA; Howard Hughes Medical Institute, University of Wisconsin-Madison, Madison, Wisconsin, 53706, USA; Department of Urology, Stanford University School of Medicine, Stanford, CA, USA; The David H. Koch Institute for Integrative Cancer Research, MIT, Cambridge, MA 02139, United States

## Abstract

Stem cell self-renewal is essential to development and tissue repair. The *C. elegans* LST-1 protein is a pivotal regulator of self-renewal and oncogenic when misexpessed. Here we define regions within the LST-1 protein that provide molecular insights into both its function and regulation. LST-1 self-renewal activity resides within a predicted disordered region that harbors two KXXL motifs. These KXXL motifs mediate LST-1 binding to FBF, a broadly conserved Pumilio/PUF RNA-binding protein that represses differentiation. Point mutations of the KXXL motifs abrogate LST-1 self-renewal activity. Therefore, FBF binding is essential to LST-1 function. A second distinct region regulates LST-1 spatial expression and primarily affects LST-1 protein turnover. Upon loss of this regulatory region, LST-1 protein distribution expands and drives formation of a larger than normal GSC pool. Thus, LST-1 promotes self-renewal as a key FBF partner, and its spatial regulation helps determine size of the GSC pool.

**IMPACT STATEMENT:** A key stem cell regulator partners with a broadly conserved PUF RNA-binding protein to drive self-renewal and maintain a stem cell pool.

## INTRODUCTION

A central paradigm in stem cell biology is that niche signaling regulates key target genes to promote self-renewal. Examples of niches and niche signaling pathways abound (e.g. Lander et al., 2012), but direct targets of niche signaling – the key genes activated in stem cells to drive self-renewal – have, for the most part, been elusive, with a handful of exceptions. One such is the *myc* gene, a target of Notch, BMP and Wnt signaling in mammalian stem cells (Moore & Lemischka, 2006). Another is the *lst-1* (<u>l</u>ateral <u>si</u>gnaling <u>t</u>arget-1) gene, a target of GLP-1/Notch signaling in nematode germline stem cells (GSCs) (**Figure 1A**) (Kershner, Shin, Hansen, & Kimble, 2014; Lee, Sorensen, Lynch, & Kimble, 2016). The LST-1 protein stands out as essential for self-renewal, and acts redundantly with another target of niche signaling, SYGL-1 (Kershner et al., 2014). LST-1 protein is normally restricted to the GSC pool region but becomes oncogenic when ubiquitously expressed (Shin et al., 2017). While the biological significance of LST-1 is unambiguous, the challenge now is to understand, in molecular terms, how LST-1 executes its key role in stem cell self-renewal and how it is regulated.

**Figure 1.**
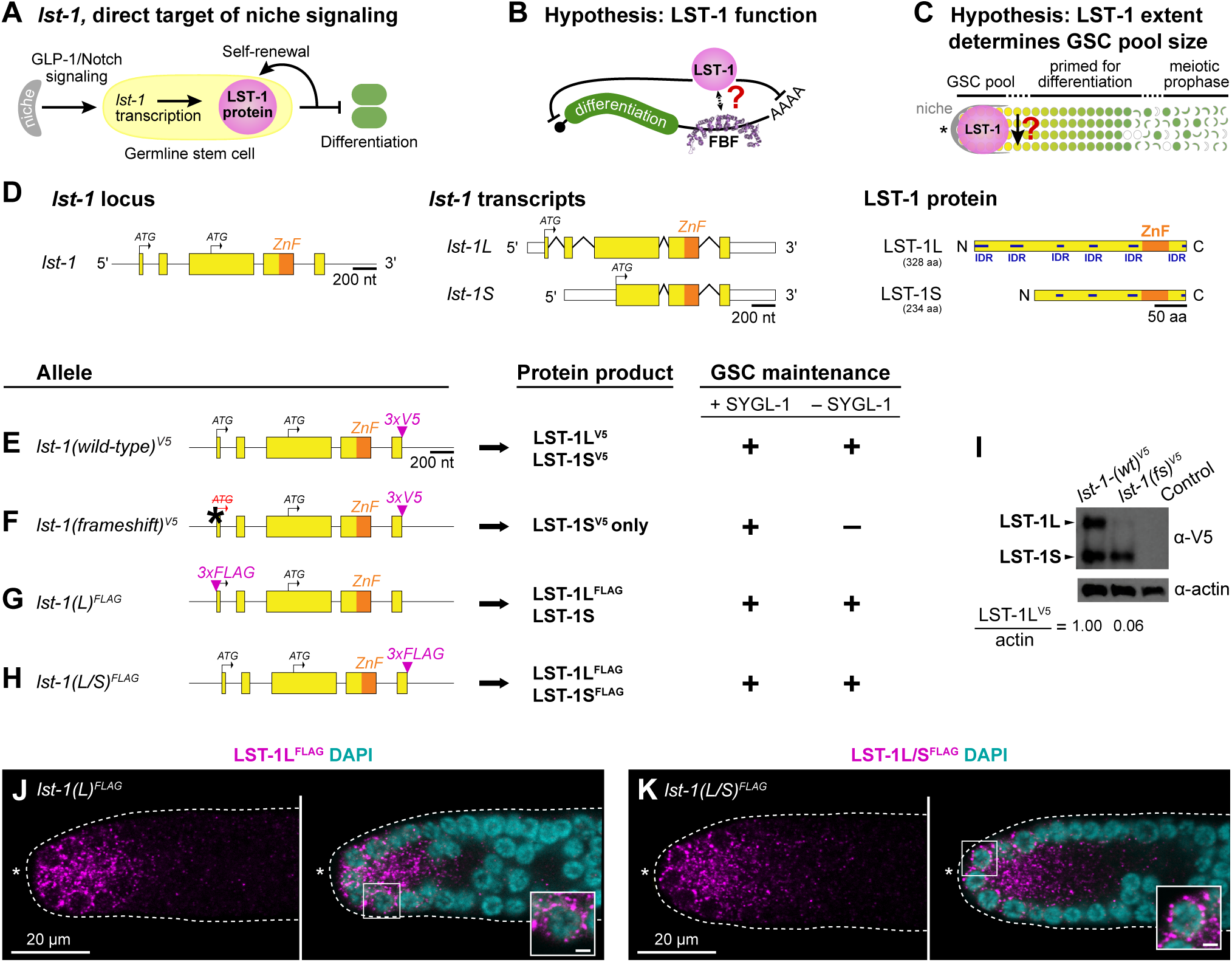
LST-1 isoforms and their role in germline stem cells. **A.** Niche signaling activates *lst-1* transcription in germline stem cells (Kershner et al. 2014; Lee et al. 2016). LST-1 protein drives self-renewal and inhibits differentiation. **B.** Hypothesis that LST-1 promotes stemness post-transcriptionally via interaction with FBF, modified from Shin et al. (2017). **C.** Hypothesis that LST-1 distribution defines extent of GSC pool in the germline tissue, modified from Shin et al. (2017). LST-1 loss (downward black arrow) launches differentiation. **D.** Left panel, the *lst-1* locus encodes two transcripts: *lst-1L(long)* and *lst-1S(short).* Each transcript (middle panel) produces a respective protein product (right panel). For locus and transcripts: exons (boxes); introns (lines between exons); open reading frame (yellow); single Nanos-related zinc finger (ZnF) (orange); untranslated regions (white boxes); start codons for each protein isoform (black arrows). For protein diagrams: coding sequence (yellow); Nanos-related zinc finger (ZnF) (orange); intrinsically disordered regions (IDR) (dark blue), predicted with DISOPRED3 (Buchan & Jones, 2019; Jones & Cozzetto, 2015). **E-H.** Left column, alleles to study LST-1L in GSC self-renewal. Conventions as described in **Figure 1D**; epitope tags (magenta); frameshift mutation (asterisk, ATG to ATΔ). Middle column, predicted protein products. Right column, GSC maintenance assay results. We assayed each allele in a *sygl-1(ø)* mutant background or by *sygl-1* RNAi. GSC maintenance was scored positive (+) if the vast majority (>90%) produced many progeny and negative (-) if all lacked GSCs. See **Supplementary Figure S1B** for more detailed information. **I.** Western blot from whole worm lysate probed with α-V5 to detect epitope-tagged LST-1 protein. LST-1L and LST-1S are visible and labeled with black arrowheads. Actin serves as a loading control, and the ratio between LST-1L and actin was calculated using Fiji/ImageJ. Control for V5 antibody specificity is wild-type N2. **J-K.** Representative single confocal Z-slices from middle plane of the distal region of an extruded gonad, stained with α-FLAG to detect the epitope-tagged LST-1 protein. N-terminal 3xFLAG epitope tags only the LST-1L isoform (**J**), while C-terminal 3xFLAG epitope tags both LST-1L and LST-1S isoforms (**K**). Dotted line delineates gonad boundary, asterisk marks distal end. Left panels, FLAG staining alone (magenta); right, merge of FLAG (magenta) and DAPI staining (cyan). Inset shows perinuclear staining; scale bar is 1µm. **E-K.** Alleles indicated are as follows: *lst-1(wildtype(wt))*^*V5*^ is *lst-1(q1004)*; *lst-1(frameshift(fs))*^*V5*^ is *lst-1(q1198)*; *lst-1(L)*^*FLAG*^ is *lst-1(q926)*; *lst-1(L/S)*^*FLAG*^ is *lst-1(q895)*.

LST-1 has been proposed to work in a macromolecular complex with PUF RNA-binding proteins, FBF-1 and FBF-2 (collectively FBF), to repress differentiation-promoting RNAs (**Figure 1B**) (Shin et al., 2017). This idea is based on several lines of evidence. First, LST-1 harbors a predicted Nanos-like zinc finger (Kershner et al., 2014) (**Figure 1D**) and the protein is cytoplasmic and granular (Shin et al., 2017), both consistent with a role in RNA regulation. Second, LST-1 interacts in yeast with FBF-1 and FBF-2 (Shin et al., 2017). Third, LST-1 cannot form tumors in the absence of FBF, suggesting that its self-renewal activity is dependent on FBF (Shin et al., 2017). Finally, LST-1 contributes to the repression of *gld-1*, an established FBF target mRNA in GSCs (J. L. Brenner & Schedl, 2016; Shin et al., 2017). While the model is attractive, it has not yet been tested *in vivo* in nematodes, nor is it known whether LST-1 must directly bind FBF to exert its self-renewal activity.

LST-1 spatial restriction to the GSC pool region suggested that the extent of *lst-1* expression might determine size of the GSC pool (**Figure 1C**) (Shin et al., 2017). Consistent with that idea, ubiquitous expression of full-length LST-1 throughout the germline drove formation of a tumor. One established regulator of *lst-1* expression in the distal germline is Notch signaling, which activates *lst-1* transcription within the niche (Kershner et al., 2014; Lee et al., 2016). However, virtually nothing was known prior to this work about how LST-1 protein is spatially restricted to the GSC pool region.

The LST-1 amino acid sequence provided few clues to the molecular basis of its self-renewal activity or its regulation (Kershner et al., 2014). Here we identify one LST-1 region sufficient for stem cell self-renewal and another required for spatial regulation. Within the self-renewal region, we find two short sequence motifs that mediate directly binding to FBF and are essential for LST-1 self-renewal activity. Within the regulatory region, we find that the Nanos-like zinc finger is central to spatial regulation: upon loss of this region, the distribution of LST-1 protein expands, leading to formation of a larger than normal GSC pool. Thus, LST-1 drives self-renewal as a key FBF partner, and its spatial regulation helps determine size of the GSC pool.

## RESULTS

### LST-1L isoform is critical for self-renewal

The *lst-1* locus encodes two transcripts, predicted to generate two proteins, a longer LST-1L and shorter LST-1S (**Figure 1D**) (Kershner et al., 2014). The LST-1L and LST-1S amino acid sequences overlap extensively and harbor multiple predicted intrinsically disordered regions (IDRs; regions with a high proportion of polar and charged amino acids and a low proportion of nonpolar amino acids (Dyson, 2016)) and a CCHC Nanos-like zinc finger (**Figure 1D**; **Supplementary Figure S1A**) (Kershner et al., 2014). To differentiate between the roles of LST-1L and LST-1S, we compared the self-renewal activities of wild-type LST-1 (**Figure 1E**) to a mutant predicted to make only LST-1S (**Figure 1F**). This mutant, termed *lst-1* (*frameshift)*, is a single base pair deletion in the *lst-1L* start codon should abolish production of LST-1L but leave LST-1S intact. For protein visualization, both *lst-1–wild-type(wt)* and *lst-1(fs)* carried a V5 epitope tag at the shared C-terminus (**Figure 1E, 1F**), which we indicate henceforth in superscript.

We assayed self-renewal activity of the *lst-1(fs)*^*V5*^ mutant and *lst-1(wt)*^*V5*^ in a *sygl-1* mutant background, where LST-1 is strictly required for GSC maintenance (Kershner et al., 2014). Previous studies showed that virtually 100% of *lst-1(wt)*^*V5*^ animals made a healthy fertile germline in the absence of SYGL-1, whereas strong loss-of-function *lst-1* mutants were 100% sterile with no GSCs (**Supplementary Figure S1B**) (Kershner et al., 2014; Shin et al., 2017). Here we confirm that *lst-1(wt)*^*V5*^ supports fertility, but find that *lst-1(fs)*^*V5*^ is 100% sterile with no GSCs in the absence of SYGL-1 (**Figure 1E, 1F, Supplementary Figure S1B**). By this simple assay, the *lst-1(fs)*^*V5*^ mutant phenotype is consistent with a strong loss-of-function. By western blot, we confirmed that LST-1L expression is nearly eliminated in *lst-1(fs)*^*V5*^, while the LST-1S isoform remains expressed (**Figure 1I**). We finally used an α-V5 antibody to stain *lst-1(wt)*^*V5*^ and *lst-1(fs)*^*V5*^ gonads and found similar spatially restricted staining in both (**Supplementary Figure S1C-F**). We conclude that LST-1L is necessary for GSC self-renewal activity, and that LST-1S not sufficient for activity despite expression in GSCs.

Consistent with its functional role in GSC self-renewal, we expected the LST-1L isoform to be expressed in GSCs. To test this idea, we introduced an epitope tag at the unique LST-1L N-terminus to visualize this isoform specifically. For this experiment, we used a FLAG tag because attempts to insert V5 at the N-terminus failed. We call the allele *lst-1(L)*^*FLAG*^ (**Figure 1G**). As a control, we generated *lst-1(L/S)*^*FLAG*^ by inserting the FLAG tag at the shared C-terminus to visualize LST-1L and LST-1S collectively (**Figure 1H**). Both N-terminal and C-terminal *lst-1*^*FLAG*^ variants maintained GSCs in the absence of SYGL-1 and therefore were functional (**Fig 1G**,**H; Supplementary Figure S1B**). Upon immunostaining, both LST-1(L)^FLAG^ and LST-1(L/S)^FLAG^ were expressed in the GSC region of the distal germline with similar subcellular localization (**Figure 1J, 1K**). We conclude that the LST-1L isoform is present in GSCs and that it harbors self-renewal activity.

### N-terminal LST-1 fragment is sufficient for stem cell maintenance

To delineate the region within LST-1 required for self-renewal activity, we generated a series of *lst-1* variant alleles (**Figure 2**). Each was introduced into the *lst-1(wt)*^*V5*^ locus (**Figure 2A**). Because all *lst-1* variants from this point carry the V5 tag, for simplicity we henceforth omit the V5 superscript in allele designations. In addition, we show only LST-1L diagrams for simplicity and because LST-1L is the primary isoform with self-renewal activity. To serve as a control, we also created an *lst-1(ø)* protein null mutant by deleting the entire open reading frame at the endogenous locus (**Figure 2B**). As expected (Kershner et al., 2014; Shin et al., 2017), all *lst-1(ø)* homozygotes maintained GSCs in the presence of SYGL-1 due to redundancy, but none maintained GSCs in the absence of SYGL-1 (**Figure 2F, Supplementary Figure S2A**).

**Figure 2.**
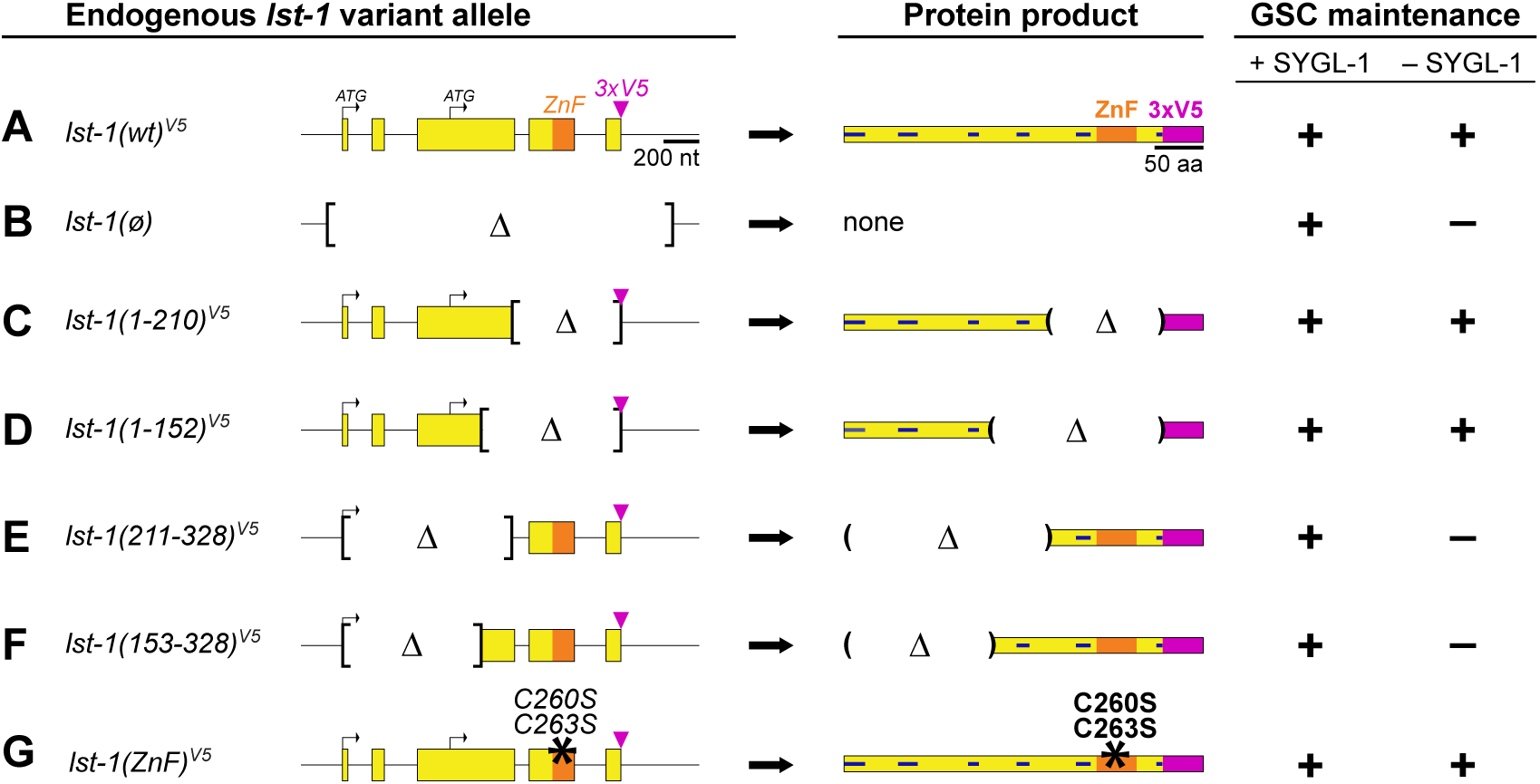
Identification of LST-1 region required for GSC maintenance. **A-G.** Left column, alleles. Only *lst-1L* is depicted for simplicity. All variants were created at the endogenous locus in *lst-1(wild-type)*^*V5*^, an otherwise wild-type allele carrying 3xV5 at its C-terminus (**Figure 1E**). Conventions as in **Figure 1D-H**; internal deletion boundaries (brackets). Middle column, predicted LST-1L protein product. Conventions as in **Figure 1D**; deletion boundaries (parentheses). For amino acid annotation of variants, see **Supplementary Figure S1A**. Right column, GSC maintenance assay and scoring as described in **Figure 1E-F**. For more detailed assay results, see **Supplementary Figure S2A**. Alleles indicated are as follows: *lst-1(wild-type)*^*V5*^ is *lst-1(q1004)*; *lst-1(ø)*, deletion allele removing the entire open reading frame as well as 139 bp upstream of the start codon and 228bp downstream of the stop codon to create a protein null (ø) is *lst-1(q869)*; *lst-1(1-210)*^*V5*^ is *lst-1(q1115)*; *lst-1(1-152)*^*V5*^ is *lst-1(q1060)*; *lst-1(211-328)*^*V5*^ is *lst-1(q1044)*; *lst-1(153-328)*^*V5*^ is *lst-1(q1119)*; *lst-1(ZnF)*^*V5*^ allele, missense mutations of two zinc finger amino acid residues (black asterisk), is *lst-1(q1032)*. These residues have been shown by others to be critical for zinc coordination and stabilization of zinc finger architecture (Hashimoto et al. 2010; Weidmann et al. 2016).

To test *lst-1* variants for function *in vivo*, each allele was assayed for GSC maintenance in the presence and absence of SYGL-1. Two variants that retained N-terminal regions, *lst-1(1-210)* and *lst-1(1-152)*, were able to maintain GSCs in the absence of SYGL-1 (**Figure 2C, 2D, Supplementary Figure S2A**). Therefore, they both harbor self-renewal activity. Complementary alleles with C-terminal parts of the protein, *lst-1(211-328)* and *lst-1(153-328)*, were unable to maintain GSCs (**Figure 2E, 2F, Supplementary Figure S2A**) (see **Figure 4** and **Supplementary Figure S5** for confirmation of germline expression). Analogous transgenic experiments performed prior to the CRISPR/Cas9 revolution gave the same results (**Supplementary Figure S2B-E**). Finally, although the truncation experiments showed that the zinc finger was dispensable for self-renewal, we explored this domain specifically by mutating two cysteine residues required for its architecture (Hashimoto et al., 2010; Weidmann et al., 2016). The *lst-1(C260S C263S)* missense mutant, dubbed *lst-1(ZnF)*, maintained GSCs in the absence of SYGL-1, like full length *lst-1(wt)* and the fragments *lst-1(1-210)* and *lst-1(1-152)*. This confirms that the zinc finger is not critical for GSC maintenance (**Figure 2G**). We conclude that LST-1 self-renewal activity resides in the N-terminal half of the protein.

**Figure 3.**
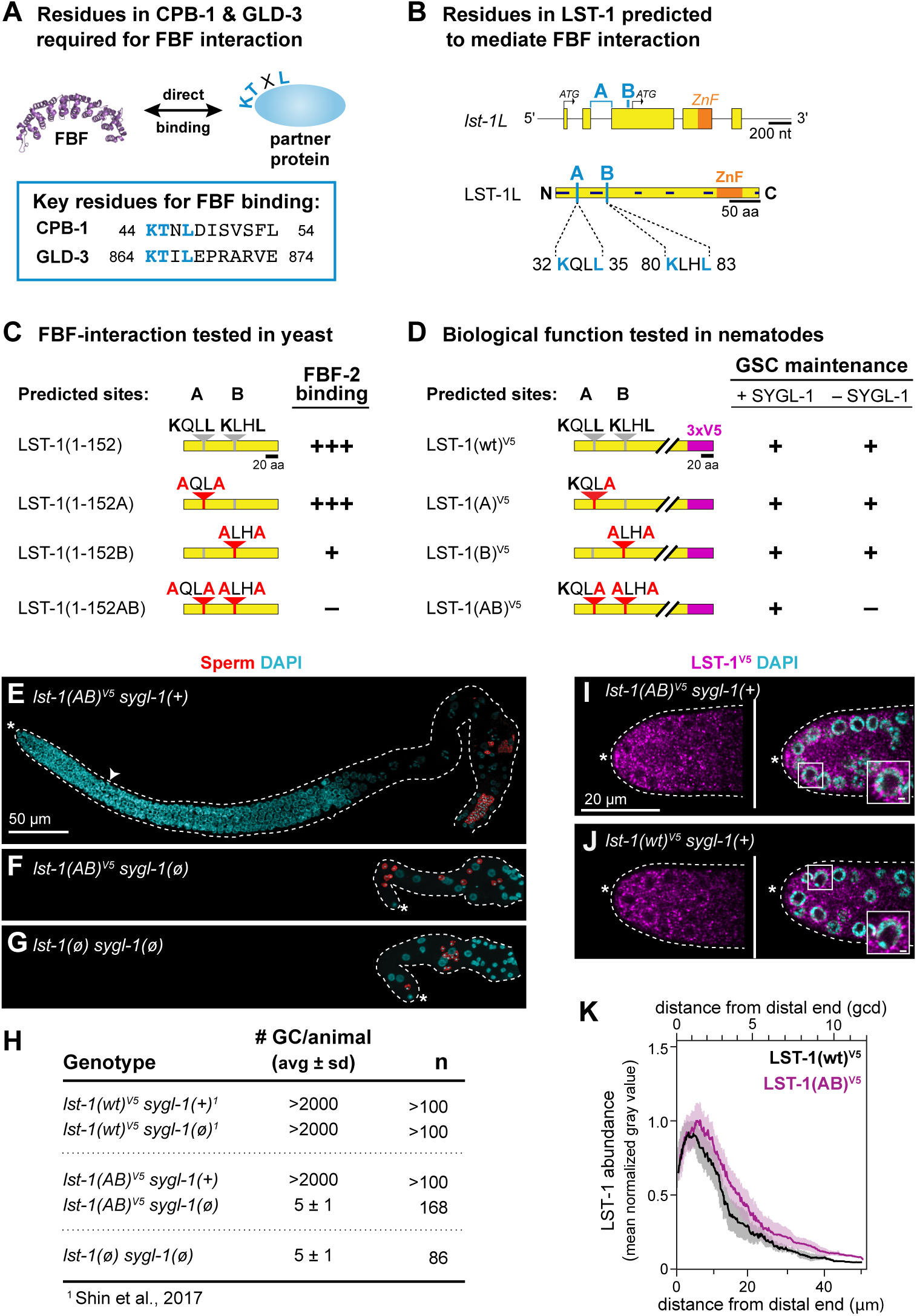
FBF-binding regions in LST-1 and their role in stem cell regulation. **A.** KTXL motif identified *in vitro* for two FBF partner proteins, CPB-1 and GLD-3 (Menichelli et al. 2013). PDB ID is 3K5Q (Wang, et al., 2009). **B.** Two predicted FBF interaction motifs, A and B, in LST-1L–specific sequence. A and B sites (light blue); other conventions as in **Figure 1D**. Top, positions of A and B in the locus. The nucleotide sequence encoding the A motif spans an intron. Below, positions of A and B in the protein (see also **Supplementary Figure S1A**). **C.** Summary of yeast two-hybrid interactions between FBF-2 and LST-1 A and B motif variants. LST-1 coding sequence (yellow); wild-type motifs (gray); mutant motifs (red); red letters indicate missense mutations. Interaction data is categorized as follows: strong binding (+++), weak binding (+), no binding (–) (see **Supplementary Figure S3** for data). **D.** Summary of *in vivo* assays testing LST-1 A and B motif variants for GSC maintenance. Protein conventions as in **Figure 1D**,**3C**. GSC maintenance assay and scoring as described in **Figure 1E-F**. For more detailed results, see **Supplementary Figure S4B**. **E-G.** Germline morphology of *lst-1(AB)*^*V5*^ allele in the presence and absence of *sygl-1*. Representative Z-projected confocal images of extruded gonads, immunostained with α-SP56 sperm antibody (red) (Ward et al. 1986) and DAPI (cyan). Scale bar in **E** is valid for all images. Conventions as in **Figure 1J-K**. **E.** *lst-1(AB)*^*V5*^ in a strain harboring wild-type SYGL-1. The full germline and progenitor zone (white carat indicates boundary) are both of normal size. **F.** *lst-1(AB)*^*V5*^ in a strain lacking SYGL-1. GSCs are not maintained: the germline is tiny and has only a few differentiated sperm (red). Other DAPI-stained nuclei belong to the somatic gonad. **G.** *lst-1(ø) sygl-1(ø)* mutants do not maintain GSCs and make only a few differentiated sperm. Previous work had characterized this phenotype with strong loss-of-function alleles (Kershner et al. 2014); here, we report the phenotype using null *(ø)* alleles. **H.** Quantitation and comparison number of germ cells (# GC) per animal. Total # GC in *lst-1(AB)*^*V5*^ *sygl-1(ø)* is indistinguishable from *lst-1(ø) sygl-1(ø)*. **I-J.** Representative single confocal Z-slices from middle plane of the distal region of an extruded gonad, stained with α-V5 to detect epitope-tagged LST-1 protein (magenta) and DAPI (cyan). Scale bar in **I** is valid for all images. Inset shows perinuclear staining; scale bar is 1µm. Conventions as in **Figure 1J-K**. **I.** LST-1(AB)^V5^ protein is restricted to distal germline, with a distribution similar to wild-type (compare to **J**). **J.** LST-1(wt)^V5^ protein is restricted to the distal germline, as previously reported (Shin et al. 2017). **K.** Quantitation of LST-1(wt)^V5^ and LST-1(AB)^V5^ protein as a function of distance from the distal end, determined by Fiji/ImageJ (see Methods for details). Lines represent the mean value of three independent replicates, each with at least 7 gonads; shading shows standard error. Sample sizes were as follows: LST-1(wt)^V5^: 3 replicates with a total of 25 germlines; LST-1(AB)^V5^: 3 replicates with a total of 36 germlines. After background from the wild-type N2 control was subtracted, LST-1(wt)^V5^ is set to 1.0 at its peak and LST-1(AB)^V5^ is normalized to this value. X-axis shows distance from distal end in microns (μm) at bottom and in germ cell diameters (gcd) at top. **D-K.** Alleles indicated are as follows: *lst-1(wt)*^*V5*^ is *lst-1(q1004)*; *lst-1(A)*^*V5*^ is *lst-1(q1124)*; *lst-1(B)*^*V5*^ is *lst-1(q1086)*; *lst-1(AB)*^*V5*^ *is lst-1(q1125)*; *lst-1(ø)* is *lst-1(q869)*; *sygl-1(ø)* is *sygl-1*(*q828)*.

**Figure 4.**
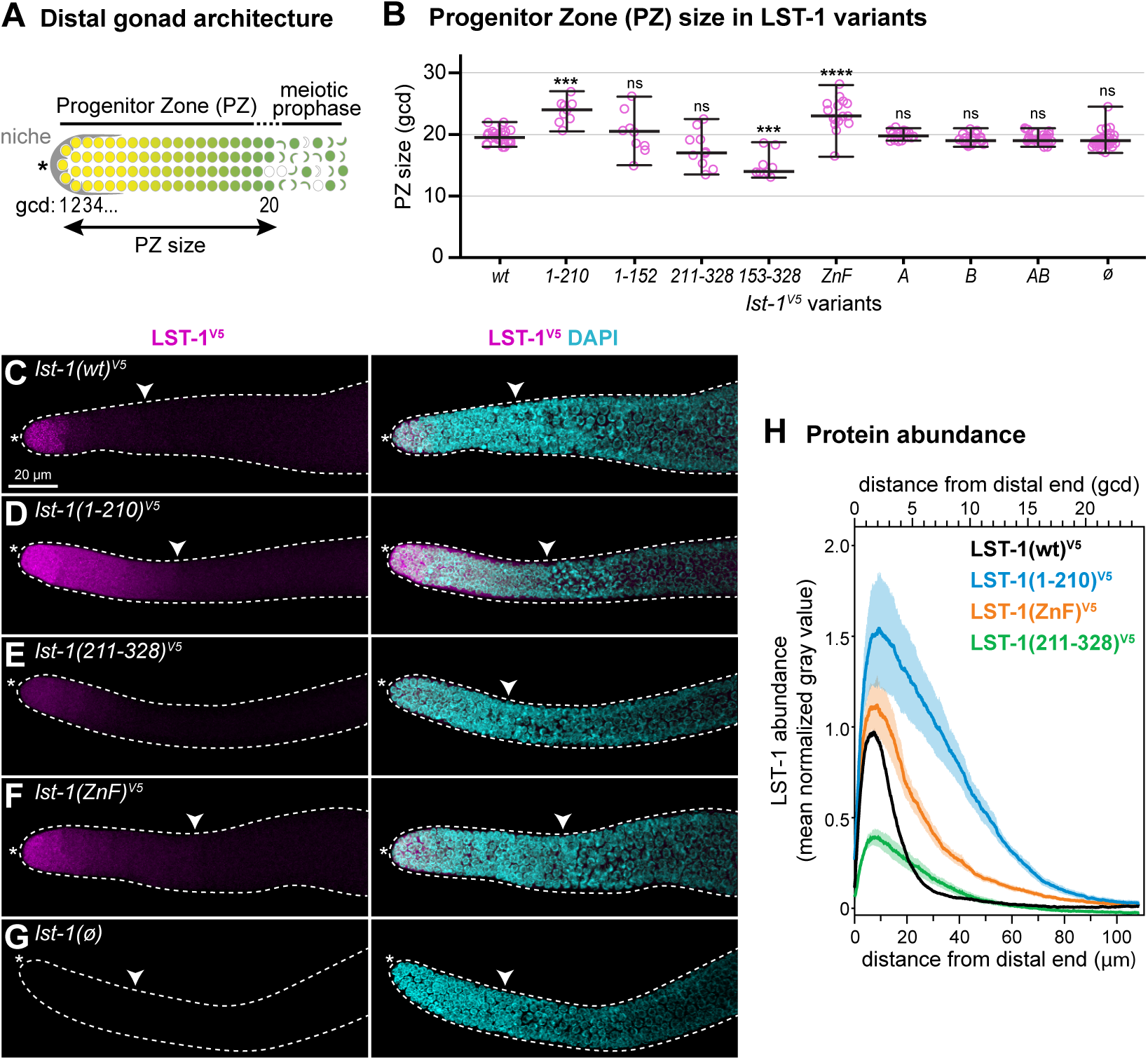
Biological readout and protein expression of LST-1 variants. **A.** Schematic of progenitor zone (PZ) in the distal gonad. Somatic niche (grey); germline stem cells in naïve state (yellow); GSC daughters primed for differentiation and transitioning toward meiotic entry (graded yellow to green); early meiotic prophase (crescent-shaped). Double-headed arrow marks PZ size, measured by convention (see Methods) in germ cell diameters (gcd) from the distal end. **B.** Progenitor zone size in *lst-1* variants measured in average gcd from the distal end. Individual data points are plotted as pink circles; middle line, median; whiskers, minimum and maximum values. n≥9 for each sample. Asterisks indicate a statistically significant difference by one-way ANOVA with Tukey’s *post hoc* test compared to *lst-1(wt)*^*V5*^: ‘****’ indicates p<0.0001, ‘***’ indicates p<0.001, ‘ns’ indicates not significant (p>0.05). **C-G.** LST-1 variant protein expression in the distal gonad. Representative confocal Z-projections of extruded gonads stained with α-V5 to detect epitope-tagged LST-1 variant protein (magenta) and DAPI (cyan). Left panel, α-V5 immunostaining; right panel, merge of α-V5 immunostaining and DAPI. Scale bar in A valid for all images. Convention as in **Figure 1J-K**; white carat marks proximal boundary of progenitor zone. **H.** Quantitation of LST-1 protein as a function of distance from the distal end, determined by Fiji/ImageJ (see Methods for details). Each line represents mean values of at least three independent replicates, each with at least 5 gonads; shading shows standard error. Total germlines analyzed were at least 23 per genotype with sample sizes as follows: LST-1(wt)^V5^: 5 replicates with a total of 41 germlines; LST-1(1-210)^V5^: 3 replicates with a total of 23 germlines; LST-1(ZnF)^V5^: 3 replicates with a total of 27 germlines; LST-1(211-328)^V5^: 3 replicates with a total of 27 germlines. After background from the *lst-1(ø)* negative control was subtracted, LST-1(wt)^V5^ was set to 1.0 at its peak and variants were normalized to this value. X-axis shows distance from distal end in microns (μm) on the bottom and in germ cell diameters (gcd) on the top. **B-H.** Alleles indicated are as follows: *lst-1(wt)*^*V5*^ is *lst-1(q1004)*; *lst-1(1-210)*^*V5*^ is *lst-1(q1115)*; *lst-1(1-152)*^*V5*^ is *lst-1(q1060)*; *lst-1(211-328)*^*V5*^ is *lst-1(q1044)*; *lst-1(153-328)*^*V5*^ is *lst-1(q1119)*; *lst-1(ZnF)*^*V5*^ is *lst-1(q1032)*; *lst-1(A)*^*V5*^ is *lst-1(q1124)*; *lst-1(B)*^*V5*^ is *lst-1(q1086)*; *lst-1(AB)*^*V5*^ *is lst-1(q1125)*; *lst-1(ø)* is *lst-1(q869)*.

Interestingly, while both *lst-1(1-210)* and *lst-1(1-152)* function properly for GSC self-renewal, these two variants differ with respect to the hermaphrodite sperm/oocyte switch: 100% of the *lst-1(1-210)* animals made the switch and were self-fertile in the absence of SYGL-1, while none of the *lst-1(1-152)* animals made the switch and instead had a sterile Mog (Masculinization of Germline) phenotype (**Supplementary Figure S2A**). A role for LST-1 in the sperm/oocyte decision was known: indeed, we note that even with SYGL-1 present, a few *lst-1(ø)* homozygotes (≤5%) failed to make the sperm-to-oocyte switch and had a Mog phenotype (**Supplementary Figure S2A)**, as previously shown for two other *lst-1* strong loss-of-function alleles (Kershner et al., 2014; Shin et al., 2017). This LST-1 effect on germline sex determination likely reflects the common molecular basis for regulation of germline self-renewal and germline sex determination as both rely on FBF RNA regulation (Sarah L. Crittenden et al., 2002; Zhang et al., 1997). Yet for the purposes of this study, we focus on LST-1 regulation of GSC maintenance via its N-terminal half.

### Two FBF binding motifs are essential for LST-1 self-renewal activity

We next sought to identify the molecular function of the N-terminal half of LST-1. LST-1 was proposed to function with FBF RNA-binding proteins to achieve self-renewal (see Introduction), two other FBF partners, CPB-1 and GLD-3, possess a consensus KTXL motif critical for FBF binding, where X is any amino acid (**Figure 3A**) (Menichelli, Wu, Campbell, Wickens, & Williamson, 2013; Wu, Campbell, Menichelli, Wickens, & Williamson, 2013). We therefore scanned the LST-1(1-152) fragment for a similar KTXL FBF-binding motif. Although no KTXL motif was found, we did find two similar sequences: KQLL (amino acids 32-35) and KLHL (amino acids 80-83). Both are conserved across Caenorhabditids (**Supplementary Figure S4A**), and we dub them A and B respectively (**Figure 3B, Supplementary Figure S1A**). Intriguingly, not only do these A and B sites reside in the N-terminal half of LST-1, they are also in the LST-1L-specific region.

We first tested A and B for FBF binding in yeast (**Supplementary Figure S3A**). Because full length, wild-type LST-1 bound to FBF-1 and FBF-2 similarly in a yeast two-hybrid assay (Shin et al., 2017), we used FBF-2 for this work. We first found that full length LST-1(wt) and LST-1(1-152) interacted similarly with FBF-2 in yeast (**Supplementary Figure S3B**) and therefore focused on LST-1(1-152) for subsequent assays. To test the importance of the A and B sites, we mutated their first and fourth positions to alanine (**Figure 3C, Supplementary Figure S3B**). Mutation of A had no appreciable effect on the yeast interaction, mutation of B reduced the interaction, and mutation of A and B in the same fragment abolished the interaction (**Figure 3C, Supplementary Figure S3B**), both in yeast growth assays (**Supplementary Figure S3C**) and β-gal assays (**Supplementary Figure S3D**). We conclude that the LST-1 self-renewal fragment has two FBF interaction motifs, and that at least in yeast, the B motif is quantitatively more important than the A motif.

To probe the biological significance of the LST-1 A and B sites in nematodes, we engineered point mutations in the two KXXL motifs at the *lst-1* endogenous locus, both individually and together (**Figure 3D**). The nematode B motif mutation was identical to that made in yeast (K80A L83A), but the nucleotide sequence encoding the A motif straddles an intron (**Figure 3B, top**), making simultaneous mutation of both K32 and L35 challenging. Because the leucine at the fourth position stood out as critical for CPB-3 and GLD-3 interactions with FBF (Menichelli et al., 2013), we chose to disrupt L35 alone to test the importance of the A motif in nematodes. We introduced the mutations into the *lst-1(wt)*^*V5*^ locus (**Figure 3D**) and generated three alleles, *lst-1(A)[L35A], lst-1(B)[K80A L83A]* and *lst-1(AB)[L35A K80A L83A]*. All were fertile and healthy when SYGL-1 was present (**Figure 3D, 3E, 3H** for *lst-1(AB)*; **Supplementary Figure S4B**).

To assay the importance of the A and B sites for GSC self-renewal, each of the three mutants was tested in the absence of SYGL-1. Both *lst-1(A)* and *lst-1(B)* maintained GSCs and were fertile (**Figure 3D, Supplementary Figure S4B**). In striking contrast, *lst-1(AB)* did not maintain GSCs: we observed a strong GSC defect characterized by self-renewal failure and the premature differentiation to sperm in an early larval stage (**Figure 3D, 3F, 3H, Supplementary Figure S4B**). This phenotype was dramatic, and indistinguishable from that of *lst-1(ø) sygl-1(ø)* (**Figure 3F, 3G, 3H**). Thus, *lst-1(AB)* has lost its self-renewal activity. Importantly, the LST-1(AB) protein was present with an abundance and distribution similar to LST-1(wt) (**Figure 3I-3K**). Therefore, LST-1(AB) protein is expressed normally, but unable to promote self-renewal. We conclude that LST-1 depends on its two KXXL FBF binding motifs for self-renewal activity.

### The C-terminal region controls spatial restriction

We next examined the *lst-1* variants for more subtle effects on GSC maintenance. To this end, we assayed size of the progenitor zone (PZ) in all of our variants (**Figure 4A**), because the PZ size may reflect the position of the regulatory network switch from stem cell state to differentiation (Sarah L. Crittenden, Leonhard, Byrd, & Kimble, 2006). PZ measurements were collected in the presence of wild-type SYGL-1 to ensure healthy germline size and organization. Most variants had a PZ size similar to wild-type (**Figure 4B**). Unexpectedly however, the PZ size was larger than wild-type for *lst-1(1-210)* and *lst-1(ZnF)* (**Figure 4B**).

We reasoned that the longer PZs in *lst-1(1-210)* and *lst-1(ZnF)* might result from an aberrant expansion of LST-1 spatial expression. To test this idea, gonads dissected from selected variants were immunostained with an α-V5 antibody; wild-type SYGL-1 was again present to ensure robust germline size and organization. LST-1(wt) protein was restricted to the same distal germline region reported previously (Shin et al., 2017) (**Figure 4C**). By contrast, LST-1(1-210) and LST-1(ZnF) were both more abundant and their expression extended more proximally than wild-type (**Figure 4D, 4F**). LST-1(211-328) was of lower abundance (**Figure 4E**). *lst-1(ø)* does not carry a V5 tag and so provided a negative control where no staining was observed (**Figure 4G**). To quantify protein abundance as a function of position within the distal gonad, we generated Z-projections (n≥5) from at least three independent experiments. We used Fiji/ImageJ to score LST-1 abundance along the gonadal axis (see Methods for details). Quantitation confirmed that LST-1(1-210) and LST-1(ZnF) proteins were more abundant and extended further proximally than LST-1(wt) (**Figure 4H**). This altered expression pattern offers a likely explanation for the enlarged PZ in the two mutants.

The increased abundance of LST-1(1-210) and LST-1(ZnF) proteins could be due to increased *lst-1* mRNA stability, increased translation or changes in LST-1 protein turnover. To query the abundance of *lst-1* mRNAs, we performed single molecule fluorescence *in situ* hybridization (smFISH) with probes designed against sequences that were intact and identical in the three variants tested but absent in the control (**Figure 5A**). The result was striking: *lst-1* RNAs in *lst-1(wt), lst-1(1-210)* and *lst-1(ZnF)* were all restricted to the distal-most 4-5 rows of germ cells and absent in the *lst-1(ø)* control (**Figure 5B-E)**. LST-1 protein expansion was therefore not due to a coordinate expansion of RNAs. Curiously, quantitation revealed that *lst-1(1-210)* and *lst-1(ZnF)* RNAs were modestly more abundant (∼30% higher) than *lst-1(wt)* RNA (**Figure 5F, 5G**), suggesting that the LST-1 zinc finger has a minor autoregulatory effect on *lst-1* mRNA abundance. Together, these quantitative analyses of *lst-1* RNA and protein indicate that the LST-1 C-terminal region mediates two distinct regulatory activities: the zinc finger downregulates *lst-1* RNA abundance, and a broader region, yet to be defined but including the zinc finger, downregulates LST-1 protein abundance and controls extent, likely through an effect on protein turnover (see Discussion).

**Figure 5.**
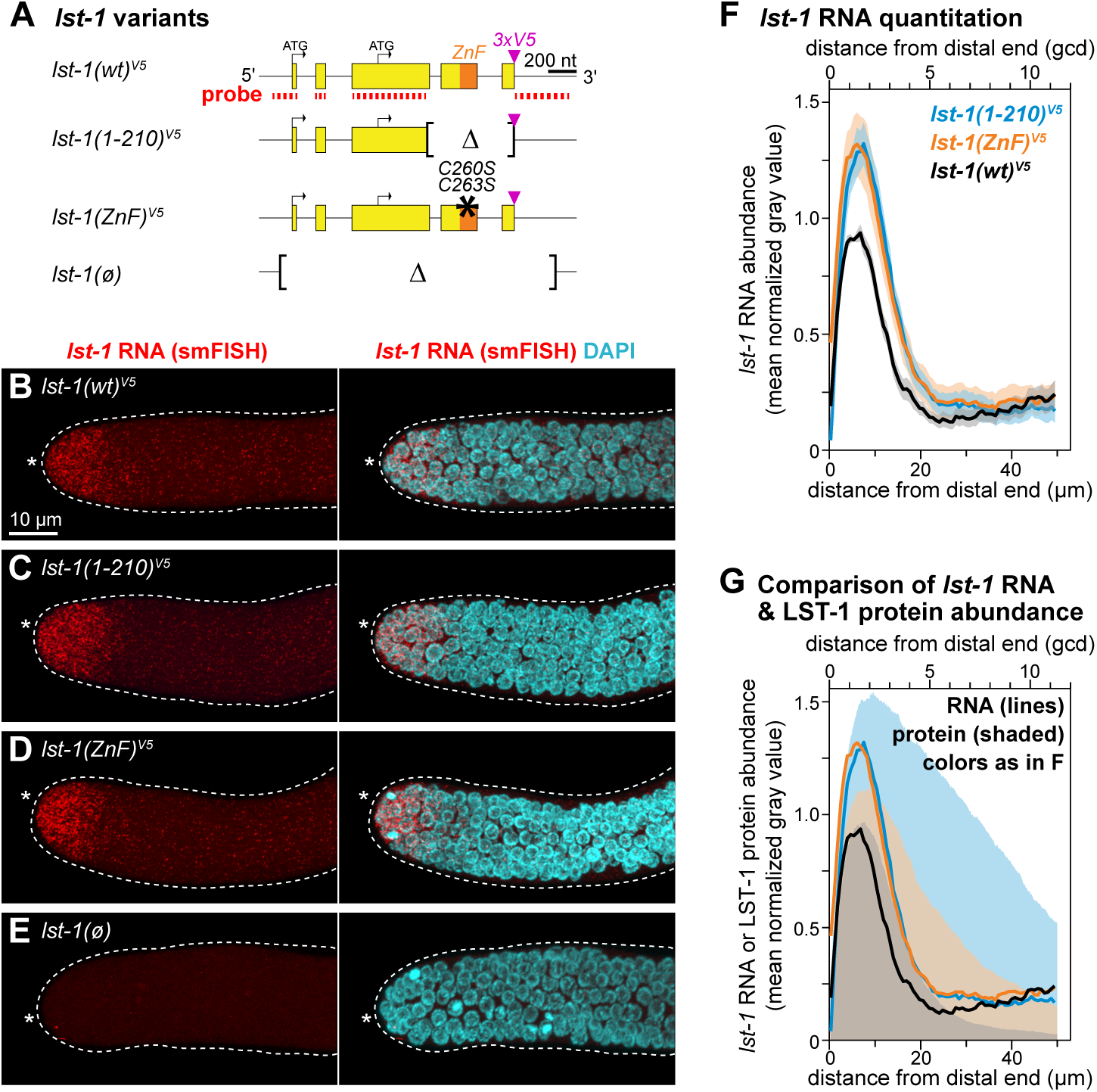
*lst-1* variant RNA expression. **A.** *lst-1* smFISH probe (red) anneals to sequences present in all strains, except the negative control *lst-1(ø). C*onventions as in **Figure 1D-H**. **B-E.** Representative Z-projected confocal images of *lst-1* RNA distribution. Extruded gonads were stained for *lst-1* RNA (red) using smFISH and DAPI (cyan). Left panel, *lst-1* RNA; right, merge of *lst-1* RNA and DAPI. Scale bar in **B** valid for all images. Conventions as in Figure **1J-K**. **F.** Quantitation of *lst-1* RNA as a function of distance from the distal end, determined by Fiji/ImageJ (see Methods for details). Each line represents mean values of three independent replicates, each with at least 9 gonads, for a total of 30 gonads analyzed per variant; shading shows standard error. After background subtraction using *lst-1(ø), lst-1(wt)*^*V5*^ was set to 1.0 at its peak and variants were normalized to this value. Distance from distal end in microns (μm) on the bottom and in germ cell diameters (gcd) on the top. **G.** Comparison of mean *lst-1* RNA values from **Figure 4F** (lines) and mean LST-1 protein values from **Figure 3F** (shading) in the distal germline. **A-G.** Alleles indicated are as follows: *lst-1(wt)*^*V5*^ is *lst-1(q1004)*; *lst-1(1-210)*^*V5*^ is *lst-1(q1115)*; *lst-1(ZnF)*^*V5*^ is *lst-1(q1032)*; *lst-1(ø)* is *lst-1(q869)*.

### Spatial extent of LST-1 determines GSC pool size

Finally, we tested the idea that the increased PZ size in *lst-1(1-210)* and *lst-1(ZnF)* mutants reflects a shift in the regulatory network from self-renewal to differentiation. To this end, we conducted *emb-30* assays (**Figure 6A**) (Cinquin, Crittenden, Morgan, & Kimble, 2010). Briefly, this assay blocks the cell cycle and stops proximal cell migration through the progenitor zone so that germ cells reveal their naïve or differentiated state *in situ* (see legend for more detail). It is the only functional assay available for GSC pool size, and we interpret the results to be a rough estimate of the number of GSCs made in each strain tested. We focused on *lst-1(wt)* and *lst-1(1-210)* for this experiment, and tested both CRISPR-induced LST-1 variants at the endogenous locus (**Figure 2A, 2C**) as well as single-copy transgenic variants inserted at a MosSCI site (**Supplementary Figure S2B, S2C**). While the LST-1 fragments assayed were identical, two critical differences existed between these experiments for historical reasons. First, the endogenous alleles were assayed in a *sygl-1(ø)* background so that GSCs were dependent on LST-1 alone, while transgenes were assayed in an *lst-1(ø) sygl-1(+)* background so that all LST-1 protein came from the transgenic allele. Second, the endogenous alleles were tagged with V5 while transgenes were tagged with HA. Remarkably, these two experiments gave virtually the same result: *lst-1(wt)* possessed an average of ∼40 germ cells in its GSC pool, whereas *lst-1(1-210)* had an average of ∼55-75 (**Figure 6B**). Thus, *lst-1(1-210)* makes a significantly larger GSC pool than *lst-1(wt)* (**Figure 6B**). Our results indicate that LST-1 extent modulates size of the GSC pool.

**Figure 6.**
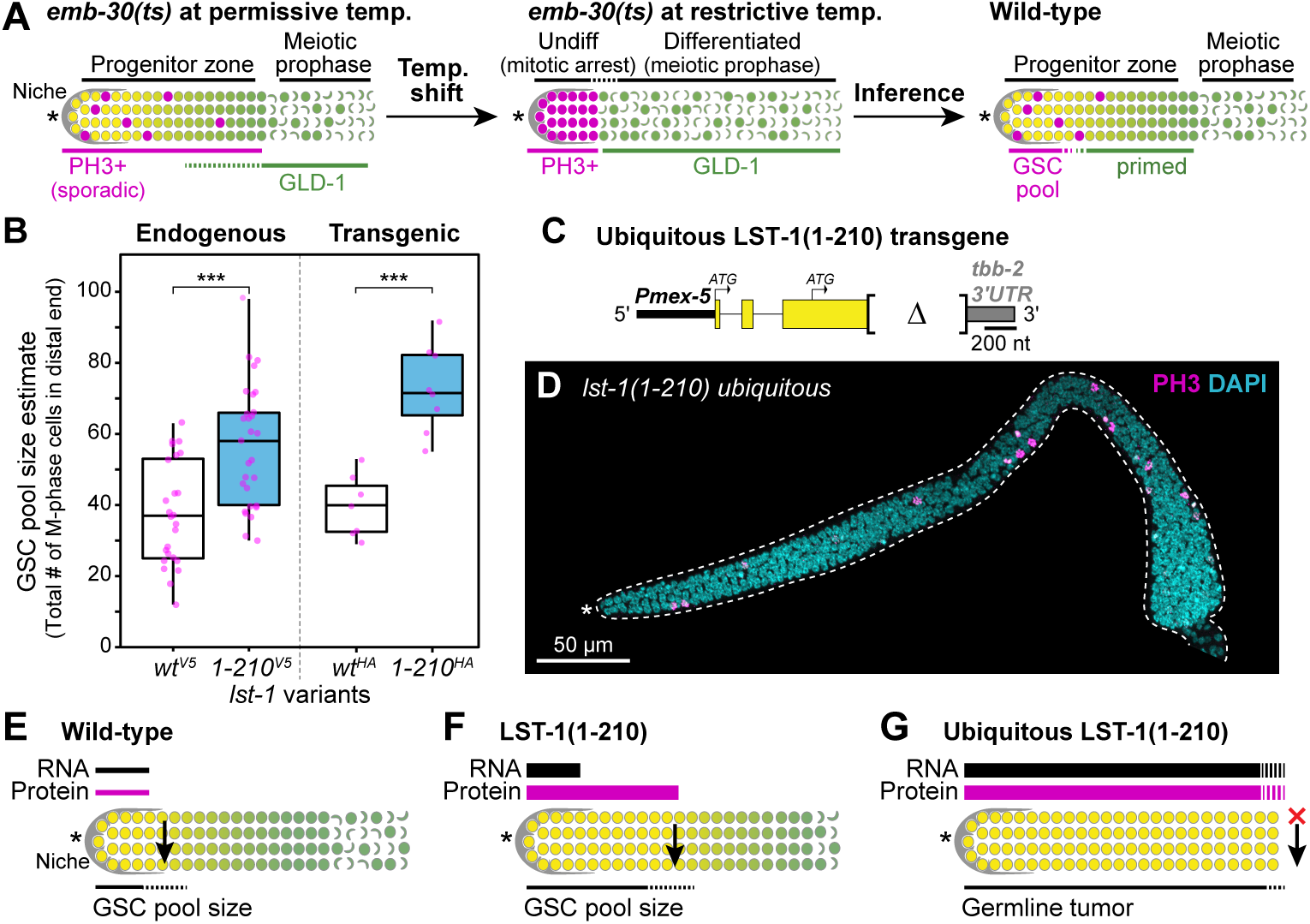
Effect of LST-1 spatial extent on GSC pool size. **A.** Schematic of *emb-30* assay to estimate GSC pool size. The *emb-30* gene encodes a component of the anaphase promoting complex (Furuta et al. 2000). In this assay, we used *emb-30(tn377ts)* to resolve two pools of cells in the PZ, one of which is inferred to be the GSC pool (Cinquin et al. 2010). Left panel: at permissive temperature, the *emb-30(ts)* progenitor zone appears normal with M-phase germ cells (pink) scattered throughout and GLD-1 levels (green) gradually increasing as germ cells move proximally towards meiotic entry. Middle panel: after shifting an *emb-30(ts)* mutant from permissive to restrictive temperature, PZ germ cells stop dividing, stop moving proximally and arrest in either of two states. Cells in the distal PZ arrest in mitotic M-phase and express the PH3 M-phase marker, but do not express the GLD-1 differentiation marker. By contrast, germ cells in the proximal PZ enter into meiotic prophase after the *emb-30(ts)* shift, and express abundant GLD-1 but no PH3. Right panel: we infer from the two pools resolved in *emb-30* gonads that the normal progenitor zone possesses a distal pool of naïve germline stem cells and a proximal pool where germ cells are in transit from the stem cell state to overt differentiation. Distal germline cartoon convention as in **Figure 4A**. **B.** Quantitation of GSC pool size, using CRISPR/Cas9-induced endogenous LST-1 variants (left) or LST-1 transgenes (right). After conducting an *emb-30* temperature shift, we visualized the distal GSC pool of cells by morphology of DAPI-stained nuclei (see Methods). *p*-values were calculated using a two-tailed *t*-test, assuming equal variance and comparing each LST-1(1-210) variant to its respective LST-1(wt) control, and ‘***’ indicates *p*<0.001. Endogenous variants were scored in a *sygl-1(ø)* background; genotypes and sample sizes are as follows: *lst-1(wt)*^*V5*^ is *lst-1(q1004) sygl-1(q828); emb-30(tn377ts)* (n=25) and *lst-1(1-210)*^*V5*^ is *lst-1(q1115) sygl-1(q828); emb-30(tn377ts)* (n=30). Transgenic variants were scored in a *sygl-1(+)* and genotypes are as follows: *lst-1(wt)*^*HA*^ is *lst-1(ok814); qSi22; emb-30(tn377ts)* (n=7) and *lst-1(1-210)*^*HA*^ is *lst-1(ok814); qSi300; emb-30(tn377ts)* (n=8). **C.** Schematic of MosSCI transgene driving ubiquitous germline expression of LST-1(1-210). Aberrant regulation was conferred with the *mex-5* promoter (black) and the *tbb-2* 3’UTR (gray). LST-1(1-210) is fused C-terminally to GGSGG linker::3xFLAG (not shown). **D.** Representative Z-projected confocal image of germline tumor driven by the single copy transgene *qSi291*[*Pmex-5*::LST-1(1-210)::GGS-GG::3xFLAG::*tbb-2* 3’ UTR] (**C**) in an *lst-1(ok814)* background to eliminate endogenous LST-1. Extruded gonad was stained with α-PH3 (magenta) to visualize cells in M-phase and DAPI (cyan). Conventions as in **Figure 1J-K**. **E-G.** Schematics illustrating the effects of LST-1(1-210) expression on GSC pool size. *lst-1* RNA (black) and LST-1 protein (magenta) expression is shown above, where line thickness correspond to quantity and line length corresponds to the extent of the expression. Below, the resultant phenotypic effects on the GSC pool size are indicated. The location of the switch from stem cell fate to differentiation, which corresponds to LST-1 protein expression, is indicated with a black arrow. Distal germline cartoon convention as in **Figure 4A**. **E.** Wild-type: *lst-1* RNA and LST-1 protein are both restricted to the distal germline and the GSC pool size is similarly restricted. **F.** LST-1(1-210): *lst-1(1-210)* RNA slightly more abundant than in wild-type, but still restricted to the distal germline (from **Figure 4**). By contrast, LST-1(1-210) protein is considerably more abundant than wild-type and expanded proximally (from **Figure 3**). GSC pool expands correspondingly (from **Figure 5**). **G.** Ubiquitous LST-1(1-210): major expansion of LST-1 leads to germline tumor formation (from **Figure 5**). The switch to differentiation fails to occur (red X).

To further interrogate the potency of the LST-1(1-210) protein for GSC maintenance, we assayed its effect when ubiquitously expressed. This assay was essentially the same as done earlier with the full length LST-1(wt) protein, which makes a massive germline tumor when placed under control of a ubiquitous germline promoter, *mex-5*, and the *tbb-2* 3’UTR (Shin et al., 2017). To ask if LST-1(1-210) might be similarly oncogenic, we made an analogous transgene, placing LST-1(1-210) under the same regulatory elements (**Figure 6C**). The strain was created and maintained with *lst-1(RNAi)* to prevent expression of the potentially oncogenic LST-1. Upon removal from *lst-1(RNAi)*, ubiquitous LST-1(1-210) drove formation of germline tumors that were composed of mitotically dividing cells, as evidenced by M-phase nuclear morphology and PH3-marked cells (**Figure 6D**). We conclude that ubiquitous LST-1(1-210) mimics LST-1(wt) in its oncogenicity. Furthermore, our data reveal that expression of LST-1(1-210) tunes the GSC pool size, and LST-1 downregulation is required to facilitate the molecular switch from stem cell state to differentiation (**Figure 6E, 6F, 6G**).

## DISCUSSION

*C. elegans* LST-1 provides a major molecular link between niche signaling and effectors of stem cell self-renewal (Kershner et al., 2014; Shin et al., 2017). Here we establish the key features of its mechanism of action at a molecular level. The LST-1 N-terminal half is responsible for self-renewal. The C-terminal half is responsible for spatial restriction of the protein in the distal germline (**Figure 7A**). Additional analyses of each region reveal critical aspects of the LST-1–FBF molecular complex and its control of stem cell fate.

**Figure 7.**
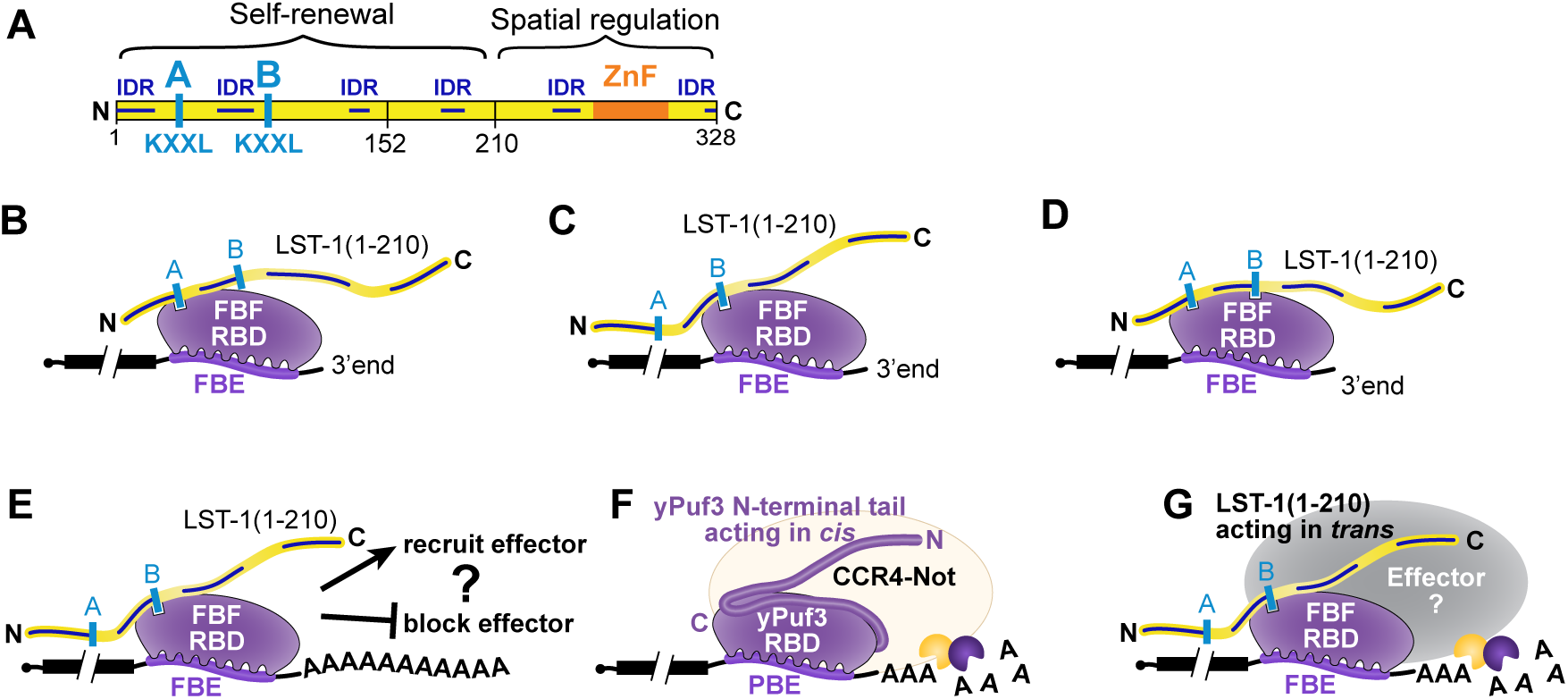
LST-1 molecular function: summary and speculation. **A.** LST-1 protein possesses one region responsible for self-renewal and another region that restricts its spatial expression. The LST-1 self-renewal region is composed largely of intrinsically disordered regions (IDR) and harbors the two KXXL motifs A and B that mediate interaction with the FBF RNA-binding protein. Conventions as in **Figure 1D, 3B**. **B-D.** Speculation about LST-1–FBF complex formation. The A and B motifs for FBF binding are functionally redundant, which means that complex formation can rely on either the A motif only (**B**) or the B motif only (**C**). Formation of these A-specific and B-specific complexes may be subject to differential regulation or offer distinct platforms for recruitment of effectors. LST-1–FBF complexes may also form via interaction with both A and B motifs (**D**), which may provide additional possibilities for regulation and function. **E-G.** Proposals for LST-1–FBF complex function. **E.** LST-1–FBF may enhance or inhibit recruitment of an effector. **F.** Possible analogy with model for yeast Puf3p, whose N-terminal tail recruits the CCR4-Not complex and promotes its deadenylase activity (modified from Webster et al. (2019). **G.** LST-1 IDRs or small linear motifs not yet identified may function in *trans* to recruit an effector.

### Dual FBF binding motifs may afford plasticity

The LST-1 N-terminal half and self-renewal region harbors two small linear motifs that mediate FBF binding, the KXXL motif. Each of these two motifs has biological activity in nematodes: LST-1 self-renewal activity remains intact when either motif is mutated, but when both are mutated, self-renewal activity is lost. The discovery of two motifs was unexpected, because other FBF partners GLD-3 and CPB-1 possess only a single KXXL motif (Campbell et al., 2012; Menichelli et al., 2013; Wu et al., 2013). The two LST-1 motifs are conserved throughout the Caenorhabditids (**Supplementary Figure S4A**), suggesting biological significance.

The existence of two KXXL motifs may afford plasticity to the LST-1–FBF complex. We do not yet know where within FBF these motifs bind but clues are available. The motifs in GLD-3 and CPB-1 interact *in vitro* at the loop between PUF repeats 7 and 8, dubbed the R7/8 loop (Menichelli et al., 2013; Wu et al., 2013). By analogy, the dual LST-1 motifs may each be able to bind the same loop. Indeed, a purified fragment harboring one LST-1 motif, called in this work the B site, binds to the R7/8 loop in the crystal structure of an FBF-2/LST-1B-site/RNA complex (C. Qiu, personal communication, May 2019). However, the LST-1 A and B motifs are unlikely to bind the same site simultaneously, raising the possibility that these dual motifs provide other opportunities. For example, some LST-1–FBF complexes may rely on the A site binding to the R7/8 loop (**Figure 7B**), while others rely on B site binding to the same loop (**Figure 7C**). This scenario introduces the possibility of considerable plasticity in the regulation and molecular configuration of the two complexes. Alternatively, the dual motifs may bind at distinct sites in FBF (**Figure 7D**). More radically, they may link two FBF proteins together, with LST-1 binding at each of their R7/8 loops. These various scenarios have important implications for configuration, stability and regulation of this critical FBF–LST-1 partnership and thus for FBF combinatorial control of RNAs and stem cell self-renewal.

### LST-1 self-renewal activity: a series of IDRs that work in *trans* with FBF

The LST-1 self-renewal region possesses two FBF-binding motifs A and B plus a series of sequences predicted to be intrinsically disordered. The LST-1(1-152) variant contains three intrinsically disordered regions (IDRs), and LST-1(1-210) has four (**Figure 7A**). Therefore, LST-1 is a *trans*-acting protein that brings a stretch of IDRs to FBF (**Figure 7E**). IDRs are commonly found in RNA-binding proteins (Calabretta & Richard, 2015) and implicated in diverse steps of RNA regulation: splicing (Chen & Moore, 2014), decapping (Jonas & Izaurralde, 2013), deadenylation (Webster, Stowell, & Passmore, 2019) and RNP granule formation (Mittag & Parker, 2018; Uversky, 2017). However, IDRs in RNA-binding proteins act within the same polypeptide and hence in *cis* with the RNA-binding domain. The LST-1 IDRs, by contrast, are not in the same polypeptide as FBF and hence work in *trans*.

The LST-1 *trans* activity could act through a range of mechanisms (**Figure 7E**). However, we suggest that LST-1 IDRs, or perhaps not-yet-identified small linear motifs interspersed within the IDRs, stabilize the formation of a complex that represses target mRNAs. Our thinking is guided by three previous studies. First, FBF binding elements regulate poly(A) tail length (Ahringer & Kimble, 1991). Second, FBF, like other PUF proteins, interacts *in vitro* with the CCR4-Not deadenylase complex (Suh et al., 2009), suggesting that FBF represses RNAs, at least in part, via deadenylation. Third, LST-1 promotes destabilization of an FBF target mRNA *in vivo* (Shin et al., 2017), suggesting that it works with FBF to promote deadenylation. Indeed, yeast Puf3 (yPuf3), a PUF RNA-binding protein from *S. cerevisiae*, possesses an N-terminal tail composed largely of IDRs and critical for interactions with the CCR4-Not complex. Remarkably, the longer the yPuf3 tail and hence larger the number of its IDRs, the greater the deadenylase activity *in vitro* (Webster et al., 2019) (**Figure 7F**). We suggest that the *trans*-acting LST-1 protein may work similarly to stabilize the interaction with an effector protein or complex (**Figure 7G**). The CCR4-Not complex is a strong candidate because of its *in vitro* interaction with FBF (Suh et al., 2009). Other possibilities exist and they are not mutually exclusive. For example, LST-1 may prevent recruitment of a positive-acting regulatory factor, such as the GLD-2/GLD-3 poly(A) polymerase (L. Wang, Eckmann, Kadyk, Wickens, & Kimble, 2002). Regardless, the discovery of an IDR-rich fragment that works in *trans* with an RNA-binding protein suggests broad, new avenues for combinatorial control.

### LST-1 downregulation and the molecular switch from stem cell state to differentiation

Spatial regulation of LST-1 in the distal germline is tight and biologically significant. Normally, *lst-1* mRNA and LST-1 protein are restricted to a germline region corresponding roughly to the GSC pool (Kershner et al., 2014; Shin et al., 2017, this work). By contrast, ubiquitous LST-1 expression drives formation of a germline tumor (Shin et al., 2017). We have found two LST-1 variants with a modest expansion in protein expression. One variant, LST-1(1-210), is a truncation fragment that lacks the C-terminal 128 amino acids; the other is a missense mutant that disrupts structural residues in the LST-1 Nanos-like zinc finger (ZnF). Both lack a functional ZnF, but LST-1(1-210) lacks additional regions as well. LST-1(1-210) and LST-1(ZnF) are more abundant and expand more proximally in the distal germline than LST-1(wt), and therefore must lack critical regions or residues controlling normal spatial regulation of the protein. As a result, we observe a delay in the switch from the stem cell state to differentiation: both variants increase the size of the progenitor zone (**Figure 4B**) and LST-1(1-210) increases size of the GSC pool (**Figure 6B, 6D**). Thus, downregulation of LST-1 protein is essential for proper cell fate determination and the switch between stem cell state and differentiation (**Figure 6E**).

The primary mechanism of LST-1 downregulation is likely via regulated protein instability. The RNAs encoding LST-1(1-210) and LST-1(ZnF) were restricted spatially as in wild-type, but the proteins were dramatically expanded (**Figure 5G**). Germ cells move proximally at a rate of about one cell per hour in the distal gonad (Rosu & Cohen-Fix, 2017), which provides a useful space-time axis. Wild-type *lst-1* RNA and LST-1 protein disappear at about the same place along this axis (Shin et al., 2017, this work), suggesting that they are both normally regulated tightly and are unstable. By contrast, LST-1(1-210) and LST-1(ZnF) proteins disappear hours later than their RNAs, with time measured by position along the axis. This change in protein turnover was particularly noticeable with LST-1(1-210). Therefore, loss of the Zn finger affects LST-1 stability, but loss of the C-terminal third is more dramatic. Earlier studies found that decreased proteasome activity leads to increased germline proliferation (Gupta et al., 2015; Macdonald, Knox, & Hansen, 2008; Mohammad et al., 2018). However, identification of the critical E3 ligase or ligases for LST-1 protein turnover remains a challenge for the future.

Regulation of protein stability as a determining factor in the fate switch from stem cell to differentiation is likely a broadly used mechanism (Werner, Manford, & Rape, 2017). Although few cases are thoroughly understood, examples exist in flies and human cells in addition to nematodes. In flies, cyclin A protein is downregulated by the Bam-dependent deubiquitinase complex to promote differentiation (Ji et al., 2017), and in human embryonic stem cells, Nanog is downregulated by ERK MAP kinase to promote differentiation (Kim et al., 2014). As more examples are uncovered, the regulation of protein stability may emerge as a ubiquitous mechanism for triggering fate switches.

### LST-1 and its role in FBF combinatorial control of stem cell regulation

This work defines LST-1 partnership with the FBF RNA-binding protein as pivotal to the LST-1 self-renewal activity. This remarkable and previously mysterious protein therefore provides an important new window into the FBF combinatorial control of stem cell regulation. Indeed, two FBF partners, LST-1 and SYGL-1, drive GSC self-renewal (Shin et al., 2017, this work). Each partner is sufficient, and at least one must be present to maintain stem cells (Kershner et al., 2014). Intriguingly, both full length SYGL-1 protein and the LST-1(1-210) self-renewal fragment consist largely of IDRs and are of comparable size. The SYGL-1 protein also possesses two KXXL motifs though their significance has not yet been tested. Nonetheless, we suggest that LST-1 and SYGL-1 are both *trans*-acting FBF partners that bring an extensive series of IDRs to their respective complexes. Analogous short, *trans*-acting RNA regulators may be more common than appreciated.

The presence of two IDR-rich FBF partners might reflect simple redundancy but also might have a more interesting role and expand the FBF repertoire for combinatorial control. They are clearly functionally redundant (Kershner et al., 2014), but in favor of individual roles, the LST-1 and SYGL-1 amino acid sequences bear no similarity to each other, and some genetic interactions differ between the two (J. L. Brenner & Schedl, 2016; Shin et al., 2017). Moreover, LST-1 localizes to perinuclear granules while SYGL-1 localizes to smaller cytoplasmic puncta, and the spatial regulation of LST-1 is much tighter than SYGL-1 (Shin et al., 2017). We therefore suggest that additional layers of regulation remain to be discovered. In this light, we note that stem cell maintenance must proceed under widely divergent physiological and environmental circumstances. Study of the factors that orchestrate responses to those circumstances, likely including LST-1 and SYGL-1, provide a tantalizing entrée to the complexities of regulation in metazoans.

## MATERIALS AND METHODS

### Nematode strains and maintenance

*C. elegans* were maintained at 20°C on Nematode Growth Medium (NGM) plates spotted with *E. coli* OP50, following established protocols (S. Brenner, 1974), except: strains containing *emb-30(tn377ts)* were maintained at 15°C; and the strain containing the *qSi291* tumor transgene was maintained on *lst-1(RNAi)* plates (see *Germline Tumor Assays* section). Wild-type was N2 Bristol strain. See **Table S1** for list of strains used in this study. We also used the balancer *LGI; LGIII hT2[qIs48]* (Siegfried & Kimble, 2002).

### CRISPR/Cas9 genome editing to generate *lst-1* alleles

See **Table S2** for list of CRISPR-induced alleles, and **Tables S4** and **S5** for additional details about their generation. We used two CRISPR/Cas9 editing methods to create alleles at the endogenous *lst-1* locus. Three alleles, *lst-1(q867), lst-1(q869) and lst-1(q926)*, were generated using a DNA-based CRISPR/Cas9 approach with a co-conversion strategy (Arribere et al., 2014; Dickinson, Ward, Reiner, & Goldstein, 2013). Briefly, the following components were microinjected into wild-type germlines: an *lst-1* sgRNA plasmid (25 ng/µl), a repair oligo designed to incorporate the desired *lst-1* mutations (500 nM) and a plasmid encoding Cas9 (*pDD162*, 50 ng/μl) (Dickinson et al., 2013) along with a *dpy-10* sgRNA (pJA58, 10 ng/µl) and repair oligo targeting the *dpy-10* locus (AF-ZF-827, 500 nM) (Arribere et al., 2014). Progeny of injected hermaphrodites were visually screened for co-injection marker editing and subsequently screened by PCR and Sanger sequencing for editing at the *lst-1* locus.

Other alleles, *lst-1(q895), lst-1(q1032), lst-1(q1044), lst-1(q1060), lst-1(q1086), lst-1(q1115), lst-1(q1119), lst-1(q1124), lst-1(q1125)* and *lst-1(q1198)*, were generated using RNA-protein complex CRISPR/Cas9 editing with a co-conversion strategy (Arribere et al., 2014; Paix, Folkmann, Rasoloson, & Seydoux, 2015). The following were microinjected into wild-type N2 (for *q895*), JK6154 (for *q1125*), JK5596 (for *q1198*) or JK5929 [*lst-1(q1004)*, which we call *lst-1(wt)*^*V5*^ for simplicity] (all other alleles): *lst-1* crRNAs (10 µM), *dpy-10* or *unc-58* co-CRISPR crRNAs (4 µM), and tracrRNA (13.6 µM) (all Alt-R™from Integrated DNA Technologies, Coralville, IA), repair oligos encoding the desired *lst-1* mutation (4 µM) and targeting the respective co-CRISPR locus (1.34 µM), and recombinant Cas9 protein (24.5 µM). Progeny of injected hermaphrodites were first visually screened for co-injection marker editing and next screened by PCR and Sanger sequencing for editing at the *lst-1* locus. All CRISPR/Cas9-generated alleles were outcrossed with wild-type at least twice prior to experimentation.

### MosSCI to generate *lst-1* transgenes

See **Table S3** for list of transgenes generated for this study, and **Table S5** for details about plasmids. The Mos1-mediated Single Copy Insertion (MosSCI) method was used to generate all transgenes (Frøkjær-Jensen, Davis, Ailion, & Jorgensen, 2012; Frøkjær-Jensen et al., 2008; Frøkjær-Jensen et al., 2014). Briefly, repair plasmids containing the gene of interest flanked by sequence targeting the *ttTi5605* insertion site were cloned using the Gibson assembly method (Gibson et al., 2009). The repair plasmids were microinjected at 50 ng/μl together with Mos1 transposase and co-injection marker plasmids into JK4950. At least three successful insertions were isolated and analyzed in our experiments, and we report one representative line in this work. During strain generation and maintenance, *lst-1*(*qSi291) [P*_*mex-5*_*::lst-1(1-210)::GGSGG linker::3xFLAG::tbb-2 3’ UTR, unc-119(+)]* and related strains were grown on *lst-1(RNAi)* feeding bacteria to prevent germline tumorigenesis (see RNAi section of Methods).

### RNA interference

RNA interference (RNAi) was performed by feeding as described (Timmons & Fire, 1998). We used *sygl-1* or *lst-1* clones from the Ahringer RNAi library (Fraser et al., 2000) and L4440 plasmid lacking a gene of interest insertion (“empty” RNAi) when an experimental control was required. HT115(DE3) bacteria cultures harboring the RNAi vectors were grown at 37°C in 2xYT media containing 25 μg/μl carbenicillin and 50 μg/μl tetracycline overnight, then were concentrated and seeded onto NGM plates containing 1mM IPTG. Bacteria were induced overnight at RT before plating worms.

### DAPI staining

To visualize nuclear morphology, we stained extruded gonads with DAPI (4′,6-diamidino-2-phenylindole) as described (S. L. Crittenden, Seidel, & Kimble, 2017), with minor modifications. Briefly, we dissected animals in PBStw (PBS + 0.1% (v/v) Tween-20) with 0.25 mM levamisole to extrude gonads, then fixed at RT for at least 15 minutes in ∼2% (w/v) paraformaldehyde diluted in PBStw. Samples were incubated overnight in −20°C methanol, washed with PBStw, then incubated with 0.5 ng/μl DAPI in PBStw to stain DNA. We mounted in either Vectashield (Vector Laboratories, Burlingame, CA) or ProLong Gold (Thermo Fisher Scientific, Waltham, MA).

### GSC maintenance and masculinization assays

For **Figures 1E-F, 2, 3D** and **Supplementary Figures S1B, S2A, S2E, S4B**, mid-L4 hermaphrodites were placed on NGM plates at 20°C. After 3-4 days, their F1 progeny were assayed for embryo production, which requires a functional germline. All fertile animals made many embryos and young larvae and were scored positive for GSC maintenance. Sterile animals were analyzed further with DAPI staining and compound microscopy. Two types of steriles were found: Mog (for Masculinization of Germline) and Glp (Germline proliferation defective). Mog germlines had a roughly normal size, harbored mitotically dividing GSCs in the distal germline, but made only sperm (no oocytes); Mogs were scored positive for GSC maintenance. Glp steriles had a very small germline made of only a few sperm. In Glp animals, we counted sperm number after DAPI staining and divided by four to estimate germ cell number. We removed *sygl-1* in some cases by feeding RNAi and in others by crossing into a *sygl-1* loss-of-function or null mutant. For RNAi, strains were plated onto *sygl-1(RNAi)* plates as mid-L4 hermaphrodites at 20°C and their F1 progeny were scored for GSC maintenance as described. In the case of Glp germlines, we quantitated the number of germ cells by DAPI staining, counting the number of mature sperm, and dividing by four (since one germ cell differentiates into four sperm).

### Progenitor zone size

Progenitor zone (PZ) size was assessed in nematodes staged to 24 hours past mid-L4 at 20°C. Extruded gonads were DAPI stained and imaged with a confocal microscope (see Microscopy). We examined nuclear morphology to determine PZ size, according to convention (Sarah L. Crittenden et al., 2006; Seidel & Kimble, 2015). Briefly, when germ cells exit the PZ and begin meiotic prophase, their nuclear morphology takes on a distinctive crescent shape (see **Figure 4A**). We selected a central focal plane in the distal gonad and then counted the number of cells along each edge of the tissue until we reached the distal-most cell with crescent morphology. We counted manually using the FIJI/ImageJ multi-point tool, calling each DAPI-stained nucleus a unique cell row. We averaged the two values from the two edges of the gonad together to determine PZ size.

### Immunostaining

We performed immunostaining of extruded gonads as described (S. L. Crittenden et al., 2017) with minor modifications. All strains (except the strain containing the *qSi291* tumor transgene, see Germline Tumor Assays section) were grown at 20°C and staged to 24 hours past mid-L4 stage, then dissected in PBStw with 0.25 mM levamisole to extrude gonads. Tissue was fixed in 2.5% (w/v) paraformaldehyde diluted in PBStw for 10 min, then permeabilized with PBStw + 0.2% (v/v) Triton-X for 10-15 minutes. Samples were blocked for at least 1 hour and not more than 4 hours in 0.5% (w/v) bovine serum albumin diluted in PBStw, except α-FLAG which was blocked in 30% (v/v) goat serum diluted in PBStw. Next, samples were incubated overnight at 4°C with primary antibodies diluted in blocking solution as follows: mouse α-FLAG® (M2, 1:1000, MilliporeSigma, St. Louis, MO), rabbit α-GLD-1 (1:100, gift from E. Goodwin), mouse α-phospho-histone H3 (Ser10) (6G3, 1:200, Cell Signaling Technology, Danvers, MA), mouse α-V5 (SV5-Pk1, 1:1000, Bio-Rad, Hercules, CA), mouse α-SP56 (1:200, Gift from Susan Strome, University of California, Santa Cruz). Secondary antibodies were diluted in blocking solution and incubated with samples for at least one hour and not more than 4 hours as follows: Alexa 488 donkey α-mouse (1:1000, Thermo Fisher Scientific), Alexa 647 goat α-rabbit (1:1000, Thermo Fisher Scientific). To visualize DNA, DAPI was included at a final concentration of 0.5–1 ng/μl during a final PBStw wash performed after secondary antibody incubation. Samples were mounted in ProLong Gold (Thermo Fisher Scientific) and allowed to cure overnight before imaging. All steps were performed at RT unless otherwise indicated.

### smFISH

Single molecule fluorescence *in situ* hybridization (smFISH) (Raj, van den Bogaard, Rifkin, van Oudenaarden, & Tyagi, 2008; Voronina, Paix, & Seydoux, 2012) was performed as described (Lee et al., 2016). Custom Stellaris FISH probes were designed using the Stellaris Probe Designer Tool (Biosearch Technologies, Inc., Petaluma, CA). The *lst-1* probe set contains 40 probes targeting the 5’UTR of *lst-1L*, the coding sequence for amino acids 1-210, and the 3’UTR. Probes were labeled with CAL Fluor Red 610 and used at a final concentration of 0.25 μM. Probe sequences are available by request.

### *emb-30* assay

The assay was performed as previously described (Cinquin et al., 2010; Shin et al., 2017) with minor modifications. Briefly, strains were maintained in a programmable incubator at 15°C until 36 hours past L4, then transitioned to 25°C for an additional 12 hours. Gonads were extruded, fixed, and stained for α-PH3, α-GLD-1 and DAPI (see Immunostaining). We imaged gonads by confocal microscopy (see Microscopy). To analyze the images, we used the DAPI channel to determine the “M-phase boundary” between presence and absence of arrested M-phase cells. In cases where a single M-phase cell was found more than three cell rows proximal to all other M-phase cells, that cell was disregarded for determining the boundary. We counted all cells distal to the M-phase boundary, including arrested M-phase cells and cells likely still arrested but not with a typical M-phase morphology, using the multipoint tool in Fiji/ImageJ (Schindelin et al., 2012). Germlines with excessively fragmented distal nuclei were excluded from the counts as cell numbers could not be determined (20-60% per experiment).

### Germline tumor assays

To induce ubiquitous expression of LST-1(1-210) using the *qSi291* tumor transgene, L4 P0 animals were transferred from *lst-1* RNAi bacteria to OP50-seeded NGM plates. Experiments were performed at 15°C to maximize tumor penetrance (Shin et al., 2017). After removal from RNAi, subsequent generations were assayed by dissection microscope and showed increasing tumor penetrance (n>100 for all): in F1, we observed no animals with tumors in both germline arms; in F2, ∼60% had tumors; in F3, ∼90% had tumors. For **Figure 6D**, we dissected and stained F3 generation L4-staged animals.

### Microscopy

All gonad images were taken using a laser scanning Leica TCS SP8 confocal microscope fitted with Photomultiplier (PMT) and Hybrid (HyD) detectors and run with LAS software version 3.3.1 or X (Wetzlar, Germany). A 63×/1.40 CS2 HC Plan Apochromat oil immersion objective was used, except for **Figure S2**, which used a 40×/1.30 CS2 HC PL APO oil immersion objective. All images were taken using the standard scanner with 400-700 Hz scanning speed and 100-300% zoom. To prepare figures, Adobe Photoshop was used to equivalently and linearly adjust contrast among samples to be compared.

### Fluorescence quantitation

Immunostaining quantitation in **Figures 3K** and **4H** was performed with Fiji/ImageJ (Schindelin et al., 2012) using images taken under identical conditions across all samples. In **Figure 3K**, we performed three independent experiments consisting of at least 7 gonads per genotype for a total of at least 21 gonads per genotype. In **Figure 4H**, we performed at least three independent experiments consisting of at least 5 gonads per genotype, with a total of at least 23 gonads per genotype. To collect intensity data from our images, we adapted a workflow from the literature (J. L. Brenner & Schedl, 2016). First, the sum of all Z-slices for each gonad was projected onto a single plane. A freehand line, 50 pixels wide and at least 80 μm long that bisected the gonad, was drawn manually starting from the distal tip of the tissue. Next, the pixel intensity data for the V5 channel along the line was obtained using the Plot Profile feature. We averaged the raw pixel intensity for at least 5 gonads at every x value to generate an average protein expression plot for each genotype in a given experiment. Next, to adjust for non-specific background staining, we subtracted the average intensity of the respective negative control at each x value from the average protein expression curves. We then normalized each average protein expression curve using the maximum and minimum values of the respective average wild type (*lst-1(wt*^*V5*^)) plot. Finally, to generate the plots shown, the adjusted (background subtracted and normalized) protein expression plots for each genotype were averaged among at least three experiments. Standard error at each x value was calculated among the three independent replicates for each genotype. The number of germ cell diameters (gcd) along the x-axis were calculated using a conversion factor of 4.4 gcd/µm (Lee et al., 2016).

smFISH quantitation in **Figure 5F** was performed similarly to the immunostaining quantitation described above, with minor modifications. After Z-projection, an average gonad-specific background level was also collected and subtracted from the raw values. This was done by using the rectangle tool to create a 2 µm square box which was manually placed on the image where no transcripts could be seen by eye. This was repeated for three separate locations along the gonad: one distally within 50 µm of the distal tip, one centrally between 50-100 µm from the distal tip, and one proximally between 100-150 µm from the distal tip. For each location, the measure feature was used to collect the average pixel intensity within the 2×2 µm box. The values obtained for each location were then averaged together to yield the final background value for the individual gonad. This gonad-specific background value was subtracted from the raw values of the respective gonad and we proceeded with quantitation as described above (*i.e.* plot profile, averaging, background subtraction, and normalizing). We performed three independent experiments consisting of at least 9 gonads per genotype for a total of 30 gonads per genotype analyzed. To compare between data sets that were collecting using different zoom factors, we condensed each average RNA expression plot by calculating a rolling average of either 4 or 5 ×- and y-values. After adjustment, the respective x-values across all data sets were essentially equal and differed by no more than 0.02 µm. For smFISH, the genotype of the negative control was *lst-1(q869)*, which harbors a deletion in the *lst-1* locus spanning from 139 bp upstream of the start codon to 228 bp downstream of the stop codon. Of note, five of the 40 smFISH probes used were predicted to anneal in the *lst-1(ϕ)* negative control.

### Yeast two-hybrid

Modified yeast two-hybrid assays were performed as described (Bartel & Fields, 1997). Briefly, LST-1 variants were amplified from cDNA and cloned into the Gal4 activation domain plasmid pACT2 using the Gibson assembly method (Gibson, 2009). We also used plasmid pJK2017, which is FBF-2 cDNA (a.a. 121– 632) fused to the LexA binding domain in the pBTM116 backbone (Shin et al., 2017). Activation and binding domain plasmid pairs were co-transformed into L40-*ura3* strain (*MATa, ura3-52, leu2-3,112, his3Δ200, trp1Δ1, ade2, LYS2::(LexA-op)*_*4*_ –*HIS3, ura3::(LexA-op)*_*8*_ *–LacZ*) using the LiOAc method (Gietz & Schiestl, 2007). *His3* reporter activity was assayed on synthetic defined medium –Leu–Trp–His plates supplemented with varying concentrations of 3-Amino-1,2,4-triazole (3-AT) (MilliporeSigma) and compared to –Leu–Trp plates as controls. We measured *LacZ* reporter activity using the Beta-Glo® Assay System following the commercially available protocols and the yeast literature (Promega, Madison, WI) (Hook, Bernstein, Zhang, & Wickens, 2005) and luminescence was quantitated using a Biotek Synergy H4 Hybrid plate reader with Gen5 software (Winooski, VT). A complete list of plasmids used in yeast-two hybrid assays is available in **Table S5**.

### Western blots

For **Figure 1I**, samples were prepared by boiling ∼50 unstaged adult worms in sample buffer (60 mM Tris pH 6.8, 25% glycerol, 2% SDS, 0.1% bromophenol blue with 700 mM beta-mercaptoethanol). For **Figure S4**, we grew yeast transformants in –Leu–Trp liquid media and prepared samples by boiling yeast in sample buffer. Subsequent analysis was conducted on a 12% SDS-PAGE gel and blots were probed with either mouse α-V5 (SV5-Pk1, 1:1000, Bio-Rad), mouse α-HA (HA.11, 1:1000, Covance, Burlington, NC) or mouse α-actin (C4, 1:40,000, MilliporeSigma) followed by donkey α-mouse horseradish peroxidase (1:10,000, Jackson ImmunoResearch, West Grove, PA). Immunoblots were developed using SuperSignal™West Pico/Femto Sensitivity substrate (Thermo Fisher Scientific) and developed using a Konica Minolta SRX-101A medical film processor (Wayne, NJ). For final figure preparations, contrast of the blot was linearly adjusted in Adobe Photoshop. For **Figure 1I**, Fiji/ImageJ was used for quantitation.

### Statistics

Where appropriate, statistical analyses are described in figure legends. Homogeneity of variance was established using Levine’s test. One-way ANOVA and Tukey’s *post hoc* tests were performed to calculate statistical significance for multiple samples. A two-tailed *t*-test assuming equal variance was performed when comparing two samples. All statistical tests were performed in R and the *p*-value cut off was 0.05.

## Supporting information

Haupt 2019 Supplement

## ACKNOWLEDGEMENTS

We thank Jonathan Doenier, Michael Green, Sarah Jayawardene, Peggy Kroll-Conner, Kimberley Law, Alex Murphy and Brandon Taylor for help generating strains central to this work, as well as Jadwiga Forster, Kyle Krueger and Charlotte Kanzler for technical assistance. We also thank members of the Kimble and Wickens laboratories for insightful discussions, and Sarah Crittenden and Brian Carrick for comments on our manuscript. We are grateful to Laura Vanderploeg and Anne Helsley-Marchbanks for assistance with figures and manuscript preparation. We thank Susan Strome (University of California, Santa Cruz) for SP56 antibodies. Some strains used in the study were provided by the *Caenorhabditis* Genetics Center, supported by the NIH Office of Research Infrastructure Programs (P40 OD010440). J.K. is an Investigator with the Howard Hughes Medical Institute; M.W. is supported by NIH R01GM050942.

## COMPETING INTERESTS

The authors declare no competing interests.

