## Supplementary material for "LST-1 acts in *trans* with a conserved RNA-binding protein to maintain stem cells": Haupt 2019 Supplement

### A LST-1 amino acid sequence

```
LST-1L
1   MCSTFILPPR RNNCNKSSST IAYSKSQHEA PKQLLQLRSE IKPLIPLNQP  50
                                      A site
                                                      LST-1S
51  SNFWESSQDS GVLSSLSSSP QQRSGLRSQK LHLTYIEKNK RVRAMIPQQQ 100
                                   B site
101 HYHAFDRPTH YNSRKTSGPP PLMRTPSSGF SSASSSENMF SGLTLSDNEN 150
     152
151 KIHEIMDPSV DVDLDMFLLP DCRYKQPVQP STSTSRNNVS QISGSSRLNG 200
               210
201 STRHVAPIVP KVAMPSELSY ANVKRSSGYV DYMPTTYNSN VTSNSSAASV 250
               C260  C263
251 PMSPGTWVRC HYCWESYVKL CQRVANLEPL ISCDGPWNWH TLYDMQGRVT 300

301 CPRLWFAQLD RAGSEMVEQM GHARNVPV                         328
```

### B Phenotype characterization

| Genotype | % Mog Sterile | % Glp Sterile | #GC in Glp animal mean ± sd | n |
|---|---|---|---|---|
| *wild-type N2* | 0 | 0 | n/a | many |
| *lst-1(ok814)*[1] | 9 | 0 | n/a | 91 |
| *lst-1(ok814) sygl-1(q828)* | 0 | **100** | 5 ± 1 | 86 |
| *lst-1(q1004)[wild-type$^{V5}$]* | 0 | 0 | n/a | 180 |
| *lst-1(q1004)[wild-type$^{V5}$] sygl-1(q828)* | 0 | 0 | n/a | 20 |
| *lst-1(q1198)[frameshift$^{V5}$]* | 7 | 0 | n/a | 130 |
| *lst-1(q1198)[frameshift$^{V5}$] sygl-1(RNAi)* | 0 | **100** | 12 ± 1 | 14 |
| *lst-1(q926)[L$^{FLAG}$]* | nd | 0 | n/a | 100 |
| *lst-1(q926)[L$^{FLAG}$] sygl-1(RNAi)* | nd | 0 | n/a | 100 |
| *lst-1(q895)[L/S$^{FLAG}$]* | nd | 0 | n/a | 100 |
| *lst-1(q895)[L/S$^{FLAG}$] sygl-1(RNAi)* | nd | 0 | n/a | 100 |

[1] Shin et al., 2017

**C** *lst-1(wild-type)$^{V5}$*

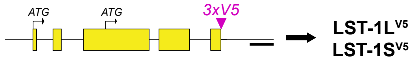


**D** *lst-1(frameshift)$^{V5}$*

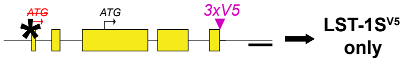


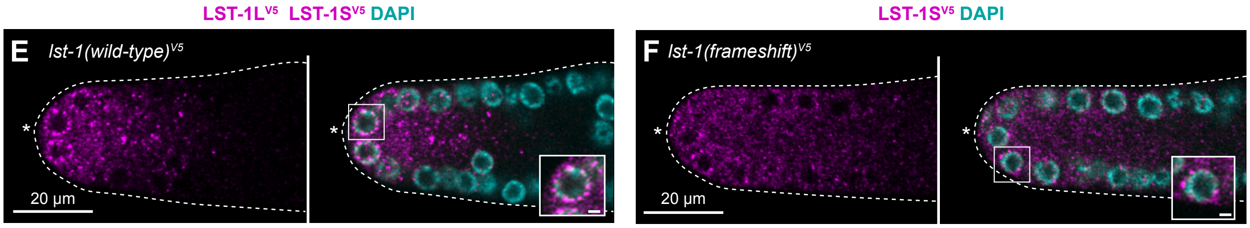


**Supplementary Figure S1. LST-1 protein sequence and L/S isoform allele characterization**
**A.** LST-1 amino acid sequence, predicted motifs and molecular basis of key *lst-1* mutants. LST-1L and LST-1S start sites marked with black arrows. Intrinsically disordered regions were predicted using DISOPRED3 (Buchan & Jones, 2019; Jones & Cozzetto, 2015) and are underlined in dark blue; putative FBF-binding sites are indicated in light blue; zinc finger underlined in orange. Residues of the boundaries for *lst-1* internal deletion variants are indicated with angled numbers, and residues altered in missense mutants are marked with asterisks.
**B.** Supplementary phenotype information for *lst-1* alleles in **Figure 1E**, **1F**, **1H** and **1I**. Mog, masculinized germline making only sperm but continuing to maintain germline stem cells. Glp, germline stem cells lost in early larval development, as in *glp-1(ø)* (Austin & Kimble, 1987). Number germ cell (# GC) per animal; normal gonads possess ~2000 germ cells (Sarah L. Crittenden et al., 2006; Kimble & White, 1981) but Glp gonads possess only 4-8 germ cells that have differentiated into sperm (Austin & Kimble, 1987). Number of GC in Glp animals obtained by counting number of sperm and dividing by four.
**C-D.** Locus diagrams of alleles to test for LST-1S germline expression and their predicted protein products (bold text). Conventions as described in **Figure 1D-H**; frameshift mutation (red asterisk, ATG to ATΔ). Genotypes are as follows: *lst-1(wild-type)$^{V5}$* allele is *lst-1(q1004)*; *lst-1(frameshift)$^{V5}$* is *lst-1(q1198)*.
**E-F.** Representative single confocal Z-slices from middle plane of the distal region of an extruded gonad, stained with α-V5 to detect the epitope-tagged LST-1 protein. Convention as in **Figure 1J-K.** Left panels, V5 staining alone (magenta); right, merge of V5 (magenta) and DAPI (cyan). Inset shows perinuclear staining; scale bar is 1µm.

### A Endogenous *lst-1* variant characterization

| Genotype | % Mog Sterile | % Glp Sterile | #GC in Glp animal mean ± sd | n |
|---|---|---|---|---|
| *lst-1(q869)[ø]* | 4 | 0 | n/a | 165 |
| *lst-1(q869)[ø] sygl-1(q828)* | 0 | **100** | 5 ± 1 | 20 |
| *lst-1(q1115)[1-210$^{V5}$]* | 0 | 0 | n/a | 106 |
| *lst-1(q1115)[1-210$^{V5}$] sygl-1(q828)* | 1 | 0 | n/a | 100 |
| *lst-1(q1060)[1-152$^{V5}$]* | nd | 0 | n/a | 100 |
| *lst-1(q1060)[1-152$^{V5}$] sygl-1(RNAi)* | **100** | 0 | n/a | 25 |
| *lst-1(q1044)[211-328$^{V5}$]* | nd | 0 | n/a | 70 |
| *lst-1(q1044)[211-328$^{V5}$] sygl-1(q828)* | 0 | **100** | 4 ± 1 | 10 |
| *lst-1(q1119)[153-328$^{V5}$]* | nd | 0 | n/a | 50 |
| *lst-1(q1119)[153-328$^{V5}$] sygl-1(RNAi)* | 0 | **100** | nd | 75 |
| *lst-1(q1032)[ZnF(C260S C263S)$^{V5}$]* | 0 | 0 | n/a | 81 |
| *lst-1(q1032)[ZnF(C260S C263S)$^{V5}$] sygl-1(q828)* | 0 | 0 | n/a | 70 |

| | Transgenic *lst-1* variant alleles | GSC maintenance + SYGL-1 | GSC maintenance – SYGL-1 |
|---|---|---|---|
| **B** | *lst-1(wt)$^{HA}$* | + | + |
| **C** | *lst-1(1-210)$^{HA}$* | + | + |
| **D** | *lst-1(1-152)$^{HA}$* | + | + |

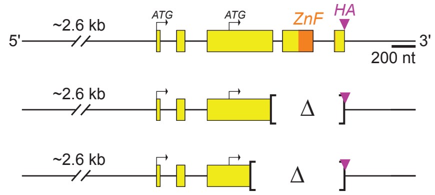


### E Transgenic *lst-1* variant characterization

| Genotype | % Mog Sterile | % Glp Sterile | #GC in Glp animal mean ± sd | n |
|---|---|---|---|---|
| *lst-1(ok814)*[1] | 9 | 0 | n/a | 91 |
| *lst-1(ok814) sygl-1(tm5040)*[2] | 0 | **100** | 6 ± 1 | 9 |
| *qSi22[lst-1(wt)$^{HA}$]; lst-1(ok814)* | 0 | 0 | n/a | 14 |
| *qSi22[lst-1(wt)$^{HA}$]; lst-1(ok814) sygl-1(tm5040)* | 0 | 0 | n/a | 14 |
| *qSi300[lst-1(1-210)$^{HA}$]; lst-1(ok814)* | 0 | 0 | n/a | 10 |
| *qSi300[lst-1(1-210)$^{HA}$]; lst-1(ok814) sygl-1(tm5040)* | 0 | 0 | n/a | 15 |
| *qSi359[lst-1(1-152)$^{HA}$]; lst-1(ok814)* | 0 | 0 | n/a | 15 |
| *qSi359[lst-1(1-152)$^{HA}$]; lst-1(ok814) sygl-1(RNAi)* | **100** | 0 | n/a | 5 |

[1] Shin et al., 2017; [2] Kershner et al., 2014

**Supplementary Figure S2. *lst-1* variant allele characterization**
**A.** Supplementary phenotype information for *lst-1* variants generated at the endogenous locus by CRISPR/Cas9 genome editing and described in **Figure 2**. Phenotype scoring convention as in **Supplementary Figure S1B**; 'nd' indicates not done.
**B-D.** Left panel: transgenic *lst-1* variant alleles, with diagram conventions as in **Figure 1D-H**. Flanking sequences upstream and downstream of the coding region were included in the transgene to preserve *lst-1* regulatory elements, and all transgenes were inserted by MosSCI at the same site on *LG II*. Alleles indicated are as follows: *lst-1(wt)$^{HA}$* is *qSi22; lst-1(1-210)$^{HA}$* is *qSi300; lst-1(1-152)$^{HA}$* is *qSi359.* Right panel: GSC maintenance assay result summary. We tested for biological activity in an *lst-1(ok814) sygl-1(tm5040)* background or *lst-1(ok814) sygl-1(RNAi)*. Assay convention as in **Figure 1E-F**.
**E.** Supplementary phenotype information for *lst-1* transgene variants in **Supplementary Figure S2B-D**, with phenotype scoring convention as in **Supplementary Figure S1B**.

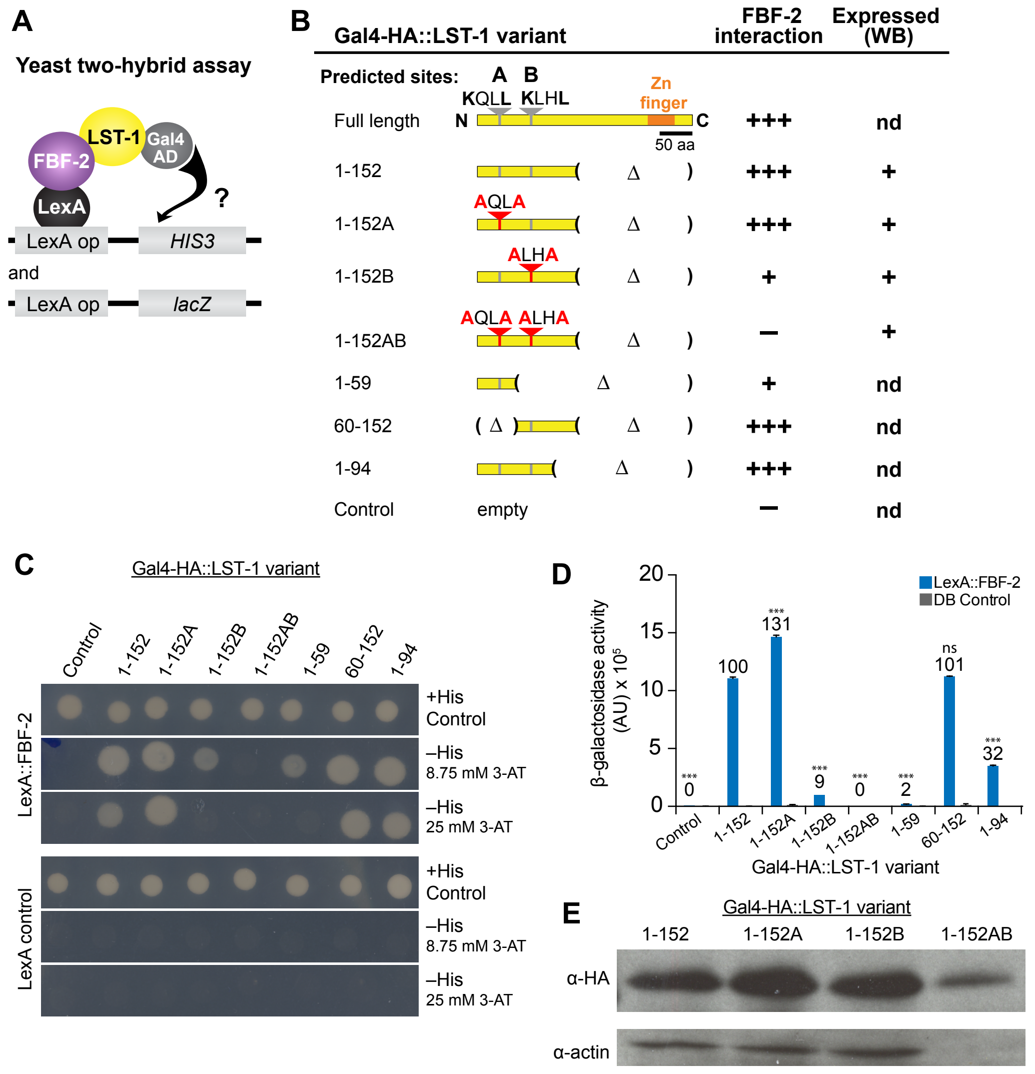


**Supplementary Figure S3. LST-1 and FBF-2 interaction assayed by yeast-two hybrid**
**A.** Yeast two-hybrid assay schematic. LST-1 variants were fused to Gal4 activation domain (AD); FBF-2(121-632), which includes all PUF repeats, was fused to LexA binding domain (BD). Interaction drives transcription of *HIS3* and *lacZ* genes.
**B.** Summary of yeast two-hybrid results. LST-1/FBF-2 interaction was measured using two assays: yeast growth (**Supplementary Figure S3C**) and β-gal activity (**Supplementary Figure S3D**). Results from both assays were consistent and are summarized by (+++) indicating robust interaction, (+) indicating weak interaction, and (–) indicating a failure to interact. For critical clones, we tested for Gal4 fusion expression by western blot (**Supplementary Figure S3E**): (+) indicates protein expression, n.d. not done. Convention for protein diagrams as in **Figure 1D**; wild-type motifs (gray); mutant motifs (red).
**C.** Yeast growth assay results. Yeast were monitored for growth on synthetic defined (SD) media with histidine as a control (+His), or lacking histidine (–His). A HIS3 competitive inhibitor (3-AT) was included in the media to improve stringency of assay.
**D.** β-gal activity assay results. Each bar is an average of two independent replicates; error bars show standard error. We indicate the activity as a percentage LST-1(1-152) with a number above each bar. Asterisks indicate a statistically significant difference by one-way ANOVA with Tukey's *post hoc* test. '***' indicates $p$<0.001, 'ns' indicates not significant ($p$>0.05).
**E.** Western blot to verify expression of select LST-1 variants in the yeast.

**A** ***Caenorhabditis* conservation of FBF interaction sites**

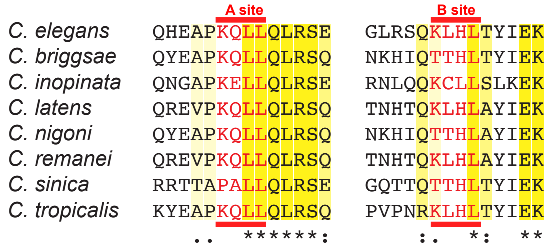


**B** **FBF interaction-related *lst-1* variant characterization**

| Genotype | % Mog Sterile | % Glp Sterile | #GC in Glp animal mean ± sd | n |
|---|---|---|---|---|
| *lst-1(q1004)[wt$^{V5}$]* | 0 | 0 | n/a | 180 |
| *lst-1(q1004)[wt$^{V5}$] sygl-1(q828)* | 0 | 0 | n/a | 20 |
| *lst-1(q1124)[A$^{V5}$]* | 0 | 0 | n/a | 243 |
| *lst-1(q1124)[A$^{V5}$] sygl-1(RNAi)* | 0 | 0 | n/a | 16 |
| *lst-1(q1086)[B$^{V5}$]* | 0 | 0 | n/a | 236 |
| *lst-1(q1086)[B$^{V5}$] sygl-1(RNAi)* | 0 | 0 | n/a | 15 |
| *lst-1(q1125)[AB$^{V5}$]* | 0 | 0 | n/a | 255 |
| *lst-1(q1125)[AB$^{V5}$] sygl-1(q828)* | 0 | **100** | 5 ± 1 | 168 |

**Supplementary Figure S4. LST-1 A/B variant allele characterization**
**A.** Amino acid sequence conservation of A and B motifs across *Caenorhabditis* species. Sequences obtained from WormBase release WS268.
**B.** Supplementary phenotype information for *lst-1* A/B site variants generated at the endogenous locus by CRISPR/Cas9 genome editing and described in **Figure 3D**. Phenotype scoring convention as in **Supplementary Figure S1B**.

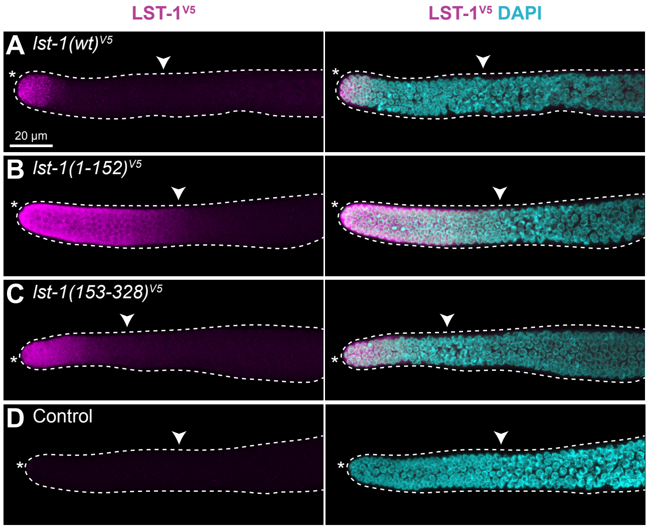


**Supplementary Figure S5. Immunostaining of additional LST-1 variants**

**A-D.** LST-1 variant protein expression in the distal gonad. Representative confocal Z-projections of extruded gonads stained with α-V5 to detect epitope-tagged LST-1 variant protein (magenta) and DAPI (cyan). Left panel, α-V5 immunostaining; right panel, merge of α-V5 immunostaining and DAPI. Scale bar in A valid for all images. Convention as in **Figure 1J-K**; white carat marks proximal boundary of progenitor zone. Alleles indicated are as follows: *lst-1(wt)*$^{V5}$ is *lst-1(q1004)*, *lst-1(1-152)*$^{V5}$ is *lst-1(q1060)*; *lst-1(153-328)*$^{V5}$ is *lst-1(q1119)*; control for V5 antibody specificity is wild-type

**S1 Table. Nematode strains used in this study**

| Name | Genotype | Reference |
|---|---|---|
| N2 | wild-type | Brenner, 1974 |
| JK4356 | lst-1(ok814) I | Singh et al., 2011 |
| JK4836 | lst-1(ok814) I; qSi22[$P_{lst-1}$::lst-1(wt)::HA::lst-1 3' end; unc-119 (+)] II | Shin et al., 2017 |
| JK4950 | lst-1(ok814) I; ttTi5605 II; unc-119(ed3) III | Shin et al., 2017 |
| JK5565 | lst-1(ok814) I; qSi291[$P_{mex-5}$::lst-1(1-210)::GGSGG linker::3xFLAG::tbb-2 3' UTR; unc-119 (+)] II | This work |
| JK5581 | lst-1(ok814) I; qSi300[$P_{lst-1}$::lst-1(1-210)::GGSGG linker::HA::lst-1 3' end; unc-119 (+)] II | This work |
| JK5583 | lst-1(q926)[3xFLAG::lst-1] I | This work |
| JK5590 | lst-1(ok814) sygl-1(q828) I/hT2[qIs48](I;III) | Shin et al., 2017 |
| JK5596 | lst-1(q867) I | This work |
| JK5622 | sygl-1(q828) I | Shin et al., 2017 |
| JK5758 | lst-1(q895)[lst-1::GGS linker::3xFLAG] I | This work |
| JK5794 | lst-1(ok814) I; qSi359[$P_{lst-1}$::lst-1(1-152)::GGSGG linker::HA::lst-1 3' end; unc-119 (+)] II | This work |
| JK5796 | lst-1(q869) I/ hT2[qIs48](I;III) | This work |
| JK5821 | lst-1(ok814) I/ hT2[qIs48](I;III); qSi22[$P_{lst-1}$::lst-1(wt)::HA::lst-1 3’ end; unc-119 (+)] II; emb-30(tn377ts) III/ hT2[qIs48](I;III) | This work |
| JK5929 | lst-1(q1004)[lst-1(wt)::3xV5] I | Shin et al., 2017 |
| JK5975 | lst-1(ok814) I/ hT2[qIs48](I;III); qSi300[$P_{lst-1}$::lst-1(1-210)::GGSGG linker::HA::lst-1 3' end; unc-119 (+)] II; emb-30(tn377ts) III/ hT2[qIs48](I;III) | This work |
| JK6011 | lst-1(q1032)[lst-1(ZnF, C260S C263S)::3xV5] I | This work |
| JK6154 | lst-1(q1086)[lst-1(B, K80A L83A)::3xV5] I | This work |
| JK6188 | lst-1(q1115)[lst-1(1-210)::GGSGG linker::3xV5] I | This work |
| JK6381 | lst-1(q1124)[lst-1(A, L35A)::3xV5] I | This work |
| JK6203 | lst-1(q1125)[lst-1(AB, L35A K80A L83A)::3xV5] I | This work |
| JK6204 | lst-1(q1125)[lst-1(AB, L35A K80A L83A)::3xV5] sygl-1(q828) I/hT2[qIs48](I;III) | This work |
| JK6254 | lst-1(q1044)[lst-1(211-328)::3xV5] I | This work |
| JK6255 | lst-1(q1060)[lst-1(1-152)::GGSGG linker::3xV5] I | This work |
| JK6256 | lst-1(q1119)[lst-1(153-328)::3xV5] I | This work |
| JK6290 | lst-1(q1044)[lst-1(211-328)::3xV5] sygl-1(q828) I/ hT2[qIs48](I;III) | This work |
| JK6319 | lst-1(q1004)[lst-1::3xV5] sygl-1(q828) I | This work |
| JK6320 | lst-1(q1115)[lst-1(1-210)::GGSGG linker::3xV5] sygl-1(q828) I | This work |
| JK6339 | lst-1(q1004)[lst-1(wt)::3xV5] sygl-1(q828) I/ hT2[qIs48](I;III); emb-30(tn377ts) III/ hT2[qIs48](I;III) | This work |
| JK6340 | lst-1(q1115)[lst-1(1-210)::GGSGG linker::3xV5] sygl-1(q828) I/hT2[qIs48](I;III); emb-30(tn377ts) III/ hT2[qIs48](I;III) | This work |
| JK6344 | lst-1(q1032)[lst-1(ZnF, C260S C263S)::3xV5] sygl-1(q828) I | This work |
| JK6399 | lst-1(q1198)[lst-1(frameshift)::3xV5] I | This work |

**S2 Table. CRISPR-induced alleles generated in this study**

| Allele | Description | Guide | Repair template | Parental strain | Method |
|---|---|---|---|---|---|
| *q867* | 1 bp deletion in *lst-1L*–specific first exon | pJK1942 | *lst-1L* frameshift repair | wild-type | **Plasmid injection co-CRISPR** (Dickinson et al., 2013) |
| *q869* | *lst-1* null mutant | pJK1965, pJK1966, pJK1967 | *lst-1* null repair | wild-type | |
| *q926* | *3xFLAG::lst-1* | pJK1942 | *3xFLAG::lst-1* repair | wild-type | |
| *q895* | *lst-1::GGS linker::3xFLAG* | *lst-1* C-term crRNA | *lst-1::GGS linker::3xFLAG* repair | wild-type | **RNA-protein complex injection co-CRISPR** (Arribere et al., 2014; Paix et al., 2015) |
| *q1032* | *lst-1(C260S C263S)::3xV5* | *lst-1* exon 4 crRNA | *lst-1(C260S C263S)* repair | JK5929 | |
| *q1044* | *lst-1(211-328)::3xV5* | *lst-1* N-term crRNA, *lst-1* exon 3 crRNA #2 | *lst-1(211-328)::3xV5* repair | JK5929 | |
| *q1060* | *lst-1(1-152)::GGSGG linker::3xV5* | *lst-1* exon 3 crRNA #1, *lst-1::3xV5* crRNA | *lst-1(1-152)::GGSGG linker:: 3xV5* repair | JK5929 | |
| *q1086* | *lst-1(K80A L83A)::3xV5* | *lst-1* exon 3 KLHL crRNA | *lst-1(K80A L83A)* repair | JK5929 | |
| *q1115* | *lst-1(1-210)::GGSGG linker::3xV5* | *lst-1* exon 3 crRNA #2, *lst-1::3xV5* crRNA | *lst-1(1-210)::GGSGG linker:: 3xV5* repair | JK5929 | |
| *q1119* | *lst-1(153-328)::3xV5* | *lst-1* N-term crRNA, *lst-1* exon 3 crRNA #1 | *lst-1(153-328)::3xV5* repair | JK5929 | |
| *q1124* | *lst-1(L35A)::3xV5* | *lst-1* exon 3 KQLL crRNA | *lst-1(L35A)* repair | JK5929 | |
| *q1125* | *lst-1(L35A K80A L83A)::3xV5* | *lst-1* exon 3 KQLL crRNA | *lst-1(L35A)* repair | JK6154 | |
| *q1198* | *lst-1(frameshift)::3xV5* | *lst-1* C-term crRNA | *lst-1::3xV5* repair | JK5596 | |

**S3 Table. MosSCI transgenes generated in this study**

| Allele | Insert description | Injected Plasmid | Parental Strain | Integration locus |
|---|---|---|---|---|
| *qSi291* | $P_{mex-5}$*::lst-1(1-210)::GGSGG linker::3xFLAG::tbb-2 3' UTR* | pJK1950 | JK4950 | *ttTi5605 II* |
| *qSi300* | $P_{lst-1}$*::lst-1(1-210)::GGSGG linker::HA tag::lst-1 3' end* | pJK1961 | JK4950 | *ttTi5605 II* |
| *qSi359* | $P_{lst-1}$*::lst-1(1-152)::GGSGG linker::HA tag::lst-1 3' end* | pJK2009 | JK4950 | *ttTi5605 II* |

**S4 Table. Oligos used to generated CRISPR alleles**

**RNA guides**

| **Name** | **Sequence (5' - 3')** | **Alleles generated** |
|---|---|---|
| *lst-1* N-term crRNA | aaaagugaacucacucuugg | *q1044*, *q1119* |
| *lst-1* exon 3 KQLL crRNA | gguugauugagcggaaucag | *q1124*, *q1125* |
| *lst-1* exon 3 KLHL crRNA | cucaggcccgaacguuguug | *q1086* |
| *lst-1* exon 3 crRNA #1 | cugauggauccaugauuucg | *q1060*, *q1119* |
| *lst-1* exon 3 crRNA #2 | aauuggugcgacaugucucg | *q1044*, *q1115* |
| *lst-1* exon 4 crRNA | ugucuccgggcacauggguu | *q1032* |
| *lst-1* C-term crRNA | uccagucuaagcaauaaaau | *q895*, *q1198* |
| *lst-1::3xV5* crRNA | uuagggauaggcuuaccgac | *q1060*, *q1115* |

**DNA repair oligos**

| **Name** | **Sequence (5' - 3')**[1] | **Alleles generated** |
|---|---|---|
| *lst-1L* frameshift repair | tccaattattcgtcggtgccatgttctttctcatttcaaaagaattatttgactattcaatcttcgcgcgagacaattgttcgacttttattcttTAtccaagagtgagttcactttttttttgaaattaaatatgtattttctctttc | *q867* |
| *lst-1* null repair | ttaaatgtttttaaagcccgcgaaattcaaaaaagctcttgcattcttgctccattgcattttcccacgcctccctccctgtttttgatccccttccctttttaataaaacataatcaactgtgcgaattatgtgaatttcaaatccc | *q869* |
| *3xFLAG::lst-1* repair | tctcatttcaaaagaattatttgactattcaatcttcgcgcgagacaatgGACTACAAAGACCATGACGGTGATTATAAAGATCATGATATCGATTACAAGGATGACGATGACAAGtgttcgacttttattcttcctccGCgagtaagttTactttttttttgaaattaaatatgtattttctctttccg | *q926* |
| *lst-1::GGS linker::3xFLAG* repair | aatggtcgaacaaatgggacacgctcgaaatgttccatcGGAGGATCCGACTACAAAGACCATGACGGTGATTATAAAGATCATGACATCGATTACAAGGATGACGATGACAAGtaagcaataaaattggtttaaatatcaattaatttatattttaccacccc | *q895* |
| *lst-1(C260S C263S)* repair | gcacagtttcacatacgactcccagGaatagtgaGaAcgaacccatgtgcccggagacattggaaccgatgctgcagac | *q1032* |
| *lst-1(211-328)::3xV5* repair | caaaagaattatttgactattcaatcttcgcgcgagacaatgaaagttgcaatgccggtgagtttaaaataaattttaagtgattttaaagaaat | *q1044* |
| *lst-1(1-152)::GGSGG linker::3xV5* repair | ctggattgacattgtcagacaatgagaacaaaatcGGAGGATCTGGAGGAggtaagcctatccctaaccctctcctcggtctagatagta | *q1060* |
| *lst-1(K80A L83A)* repair | tccagcttgtcatcttctccccaGcaGcgCtcCggcctgagaagtcaaGCTttgcatGCCacgtatatagagaagaacaagagagttcgt | *q1086* |
| *lst-1(1-210)::GGSGG linker::3xV5* repair | tctaaatggatcTacgagacatgtcgcaccaattgttccaGGAGGATCTGGAGGAggtaagcctatccctaaccctctcctcggtctagatagta | *q1115* |

| | | |
|---|---|---|
| *lst-1(153-328)::3xV5* repair | caaaagaattatttgactattcaatcttcgcgcgagacaatgcacgaaatcatggatccatcagttgacgtggatctt | *q1119* |
| *lst-1(L35A)* repair | agacgtaattaataaaaattaatttcagctgGCCcaactccgatctgaaatcaagccgctAatCccActTaatcaaccttccaatttttg | *q1124*, *q1125* |
| *lst-1::3xV5* repair | caaatgggacacgctcgaaatgttccagtcGGTAAGCCTATCCCTAACCCTCTCCTCGGTCTAGATAGTACTGGAAAGCCAATCCCAAACCCACTCCTCGGACTTGATAGCACCGGTAAGCCTATCCCTAACCCACTCCTCGGACTTGATAGCACCtaagcaataaaattggtttaaatatcaattaatttatattttac | *q1198* |

[1] Uppercase letters denote mutations (including insertions, PAM mutations and/or seed sequence mutations)

**S5 Table. Plasmids generated in this study**

**CRISPR reagents**

| **Plasmid** | **Insert Description** | **Cloning site** | **Vector Backbone** |
|---|---|---|---|
| pJK1942 | Sequence targeting *lst-1* locus (5' aaagtgaactcactcttgg 3') joined with sgRNA scaffold from pDD162 (Dickinson et al., 2013) | *Xma* I | pUC19 |
| pJK1965 | Sequence targeting *lst-1* locus (5' tattttcccacgcctggggg 3') joined with sgRNA scaffold from pDD162 (Dickinson et al., 2013) | *Xma* I | pUC19 |
| pJK1966 | Sequence targeting *lst-1* locus (5' ggatcaaaaaacagggatgg 3') joined with sgRNA scaffold from pDD162 (Dickinson et al., 2013) | *Xma* I | pUC19 |
| pJK1967 | Sequence targeting *lst-1* locus (5' gtgtgtggcaggagctaata 3') joined with sgRNA scaffold from pDD162 (Dickinson et al., 2013) | *Xma* I | pUC19 |

**MosSCI reagents**

| **Plasmid** | **Insert Description** | **Cloning site** | **Vector Backbone** |
|---|---|---|---|
| pJK1950 | $P_{mex\text{-}5}$*::LST-1(1-210)::GGSGG linker::3xFLAG::tbb-2* 3'UTR | *Spe* I | pCFJ151 |
| pJK1961 | $P_{lst\text{-}1}$ *(~2.6kb) ::lst-1(1-210)::GGSGG::HA::lst-1 3' end (~0.6kb)* | *Spe* I | pCFJ151 |
| pJK2009 | $P_{lst\text{-}1}$*(~2.6kb)::lst-1(1-152)::GGSGG::HA::lst-1 3' end (~0.6kb)* | *Spe* I | pCFJ151 |

**Yeast reagents**

| **Plasmid** | **Insert Description** | **Cloning site** | **Vector Backbone** |
|---|---|---|---|
| pJK2039 | *lst-1(1-59)* cDNA | *Xho* I, *Nco* I | pACT2 |
| pJK2040 | *lst-1(60-152)* cDNA | *Xho* I, *Nco* I | pACT2 |
| pJK2041 | *lst-1(1-152)* cDNA | *Xho* I, *Nco* I | pACT2 |
| pJK2042 | *lst-1(1-94)* cDNA | *Xho* I, *Nco* I | pACT2 |
| pJK2043 | *lst-1(1-152, K32A L35A)* cDNA | *Xho* I, *Nco* I | pACT2 |
| pJK2044 | *lst-1(1-152, K80A L83A)* cDNA | *Xho* I, *Nco* I | pACT2 |
| pJK2045 | *lst-1(1-152, K32A L35A K80A L83A)* cDNA | *Xho* I, *Nco* I | pACT2 |
